# MDR1 Drug Efflux Pump Promotes Intrinsic and Acquired Resistance to PROTACs in Cancer Cells

**DOI:** 10.1101/2021.12.02.470920

**Authors:** Alison M. Kurimchak, Carlos Herrera-Montávez, Sara Montserrat, Daniela Araiza, Jianping Hu, Jian Jin, James S. Duncan

## Abstract

PROTACs (Proteolysis-Targeting Chimeras) represent a promising new class of drugs that selectively degrade proteins of interest from cells. PROTACs targeting oncogenes are avidly being explored for cancer therapies, with several currently in clinical trials. Drug resistance represents a significant challenge in cancer therapies, and the mechanism by which cancer cells acquire resistance to PROTACs remains poorly understood. Using proteomics, we discovered acquired and intrinsic resistance to PROTACs in cancer cells can be mediated by upregulation of the drug efflux pump MDR1. PROTAC-resistant cells could be re-sensitized to PROTACs through co-administering MDR1 inhibitors. Notably, co-treatment of MDR1-overexpressing colorectal cancer cells with MEK1/2 or KRAS^G12C^ degraders and the dual ErbB receptor/MDR1 inhibitor lapatinib exhibited potent drug synergy due to simultaneous blockade of MDR1 and ErbB receptor activity. Together, our findings suggest that concurrent blockade of MDR1 will likely be required in combination with PROTACs to achieve durable protein degradation and therapeutic response in cancer.

## INTRODUCTION

PROTACs (Proteolysis-Targeting Chimeras) have emerged as a revolutionary new class of drugs that utilize the cancer cells’ own protein destruction machinery to selectively degrade essential tumor drivers (*1*). PROTACs are small molecules with two functional ends, a small-molecule end that binds to the protein of interest and the other end that binds to an E3 ubiquitin ligase (*2, 3*). The PROTAC component recruits the ubiquitin ligase to the target protein, leading to its ubiquitination and subsequent degradation by the proteasome. Benefits of PROTACs include development of drugs against previously undruggable drug targets, non-reliance on catalytic activity for degradation, as well as do not require high affinity for the drug target to achieve protein degradation (*4*). Additionally, low doses of PROTACs can be highly effective at inducing degradation, which can reduce off-target toxicity associated with high-dosing of traditional inhibitors (*3*). PROTACs have been developed for a variety of cancer targets including oncogenic kinases (*5*), epigenetic targets (*6*) and recently KRAS^G12C^ proteins (*7*). PROTACs targeting the androgen receptor or estrogen receptor are avidly being evaluated in clinical trials for prostate (NCT03888612) or breast cancers (NCT04072952) respectively.

Drug resistance represents a significant therapeutic challenge for the treatment of cancer (*8*). Resistance to PROTACs has been shown to involve genomic alterations in the core components of the E3 ligase components, such as downregulation of expression of CRBN, VHL or CUL2 proteins required for protein degradation (*9-11*). Upregulation of drug efflux pump ABCB1 (MDR1), a member of the superfamily of ATP-binding cassette (ABC) transporters has been shown to convey drug resistance to many anti-cancer drugs including chemotherapy agents, kinase inhibitors, and other targeted agents (*12*). Recently, PROTACs have been shown to be substrates for MDR1 (*10, 13*), suggesting drug efflux may represent a potential limitation for degrader therapies. Here, using BET protein and CDK9 degraders as a proof-of-concept, we applied proteomics to define acquired resistance mechanisms to PROTAC therapies in cancer cells following chronic exposure. Our study revealed a role for the drug efflux pump MDR1 in both acquired and intrinsic resistance to protein degraders in cancer cells and supports combination therapies involving PROTACs and MDR1 inhibitors to achieve durable protein degradation and therapeutic responses.

## RESULTS

### Proteomics characterization of degrader-resistant cells reveals common upregulation of the multidrug resistance protein MDR1

To explore resistance mechanisms to PROTAC therapies, we chronically exposed the ovarian cancer cell line A1847 to BET bromodomain (BD) or CDK9 degraders and carried out single-run proteomics using LC-MS/MS (*14*) comparing parental and degrader-resistant cells (**Fig. 1A**). Changes in protein abundance following chronic degrader-treatment were measured using Label-Free Quantitation (LFQ) (*15*). We generated A1847 BD or CDK9 degrader-resistant cells through chronic exposure to increasing doses of either dBET6 (*16*), MZ1 (*17*), or Thal SNS 032 (*18*). The chronically exposed A1847 cells were more resistant to BET bromodomain or CDK9 degraders than treatment-naïve (i.e., parental) cells, whereby they showed a rightward shift in dose-response cell viability curves (**Fig. 1B-C, S1A**). In contrast to parental cells, treatment of chronically exposed cells with increasing doses of BET protein degraders was insufficient to degrade BRD2, BRD3 or BRD4 and reduce BET protein target FOSL1 protein levels to extent observed in parental cells (**Fig. 1D, S1B**). Similarly, treatment of A1847 cells with increasing doses of CDK9-degrader Thal SNS 032 did not inhibit cell viability or reduce CDK9 protein levels or CDK9-mediated phosphorylation of RNA polymerase (S2) to the degree observed in parental cells, demonstrating chronic exposure to degraders reduced PROTAC degradation efficiency (**Fig. 1E**).

**Figure 1.**
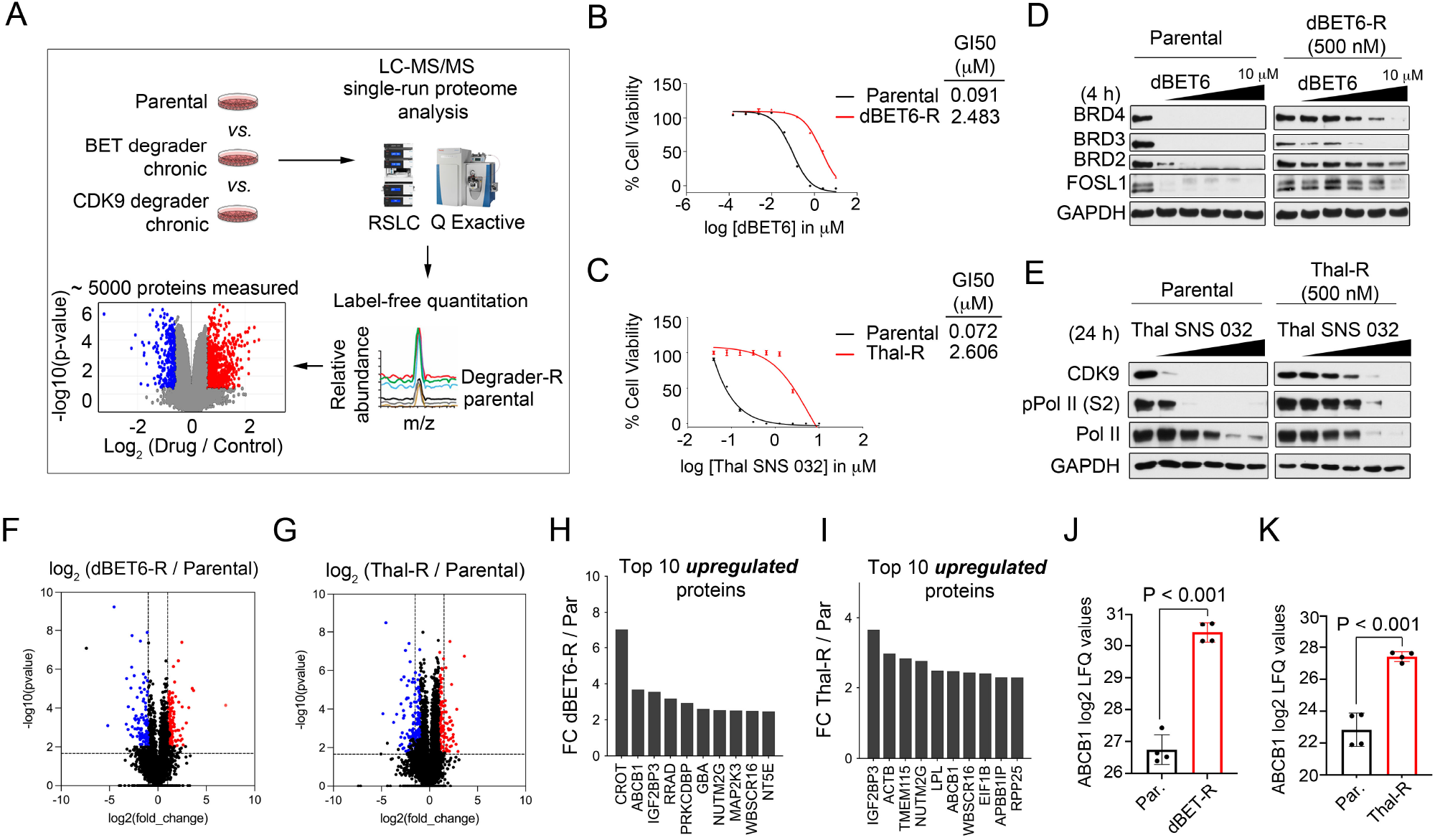
Proteomics Characterization of Degrader-Resistant Cancer Cell Lines. (A) Workflow for identifying protein targets upregulated in degrader-resistant cancer cells. Single-run proteome analysis was performed and changes in protein levels amongst parent and resistant cells determined by label-free quantitation. (B-C) A1847 cells acquire resistance to dBET6 or Thal SNS 032. Parental and dBET6 or Thal SNS 032-resistant cells were treated with escalating doses of dBET6 (B) or Thal SNS 032 (C) for 5 d and cell viability assessed by CellTiter-Glo. Degrader-R treated cell viabilities normalized to DMSO treated degrader-R cells. (D-E) Escalating doses of degraders fails to promote degradation of protein target in degrader-resistant cells. A1847 parental, dBET6-R (D) or Thal-R (E) were treated with escalating doses of dBET6 (0, 0.123, 0.370, 1.1, 3.3, or 10 μM) or Thal SNS 032 (0, 0.123, 0.370, 1.1, 3.3, or 10 μM) for 24 h and degrader targets and downstream signaling determined by western blot. Blots are representative of 3 independent blots. (F-G) Volcano plot depicts proteins elevated or reduced in dBET6-R (F) or Thal-R (G) relative to parental A1847 cells. Differences in protein log2 LFQ intensities amongst degrader-resistant and parental cells were determined by paired *t*-test Benjamini-Hochberg adjusted *P* values at FDR of <0.05 using Perseus software. (H-I) Top 10 upregulated proteins in dBET6-R (H) or Thal-R (I) relative to parental A1847 cells. (J-K) Bar graph depicts ABCB1 log2 LFQ values comparing dBET6-R (J) or Thal-R (K) relative to parental A1847 cells. Differences in ABCB1 log2 LFQ intensities amongst degrader-resistant and parental cells were determined by paired *t*-test Benjamini-Hochberg adjusted *P* values at FDR of <0.05 using Perseus software. Data present in (B), (C) are triplicate experiments SD. *p ≤0.05 by student’s t-test. Also see Figure S1, and Data File S1.

Volcano plot analysis of changes in protein abundance comparing parent and degrader-resistant cells showed significant remodeling of the proteome upon continuous exposure to BET bromodomain or CDK9 degraders (**Fig. 1F-G, S1C, Data File S1**). A comparison of the top 10 upregulated proteins in dBET6, MZ1 and Thal SNS 032 resistant cells relative to parental cells revealed 2 proteins were commonly induced, the ATP-dependent drug efflux pump, ATP Binding Cassette Subfamily B Member 1 (ABCB1) (*19*), and the RNA binding factor Insulin-Like Growth Factor 2 MRNA-Binding Protein 3 (IGF2BP3) (*20*) (**Fig. 1H-I, S1D**). Notably, ABCB1 (MDR1) is a member of the superfamily of ATP-binding cassette (ABC) transporters involved in translocation of drugs and phospholipids across the membrane and has established functions in drug resistance (*12*). MDR1 protein levels were upregulated ∼3.5 fold in dBET6-R, ∼5-5-fold in MZ1-R and ∼2.5-fold in Thal-R cells relative to parental cell lines by LFQ analysis (**Fig. 1J-K, S1E**). Similarly, chronic exposure of the breast cancer cell line SUM159 with MZ1 resulted in degrader resistance (**Fig.S1F-G**) and proteomics analysis of MZ1-resistant SUM159 cells revealed MDR1 was amongst the top 10 upregulated proteins, with an increase of ∼4.5-fold in MZ1-R cells compared to parental cells (**Fig. S1H-J, Data File S1**).

Elevated *ABCB1* mRNA and protein levels were confirmed in degrader-resistant A1847 and SUM159 cells by RT-PCR (**Fig. 2A, S2A**), immunoblot (**Fig. 2B, S2B**) and immunofluorescence (**Fig. 2C-E**), where MDR1 protein was detected at the membrane of degrader-resistant cells. Increased MDR1 drug efflux activity was detected in BET bromodomain or CDK9 degrader-resistant cells relative to parental cells using the Rhodamine 123 efflux assay (*21*) (**Fig. 2F**). Together, these findings demonstrate chronic exposure of cancer cells to BET protein or CDK9 degraders can result in increased MDR1 protein levels and drug efflux activity.

**Figure 2.**
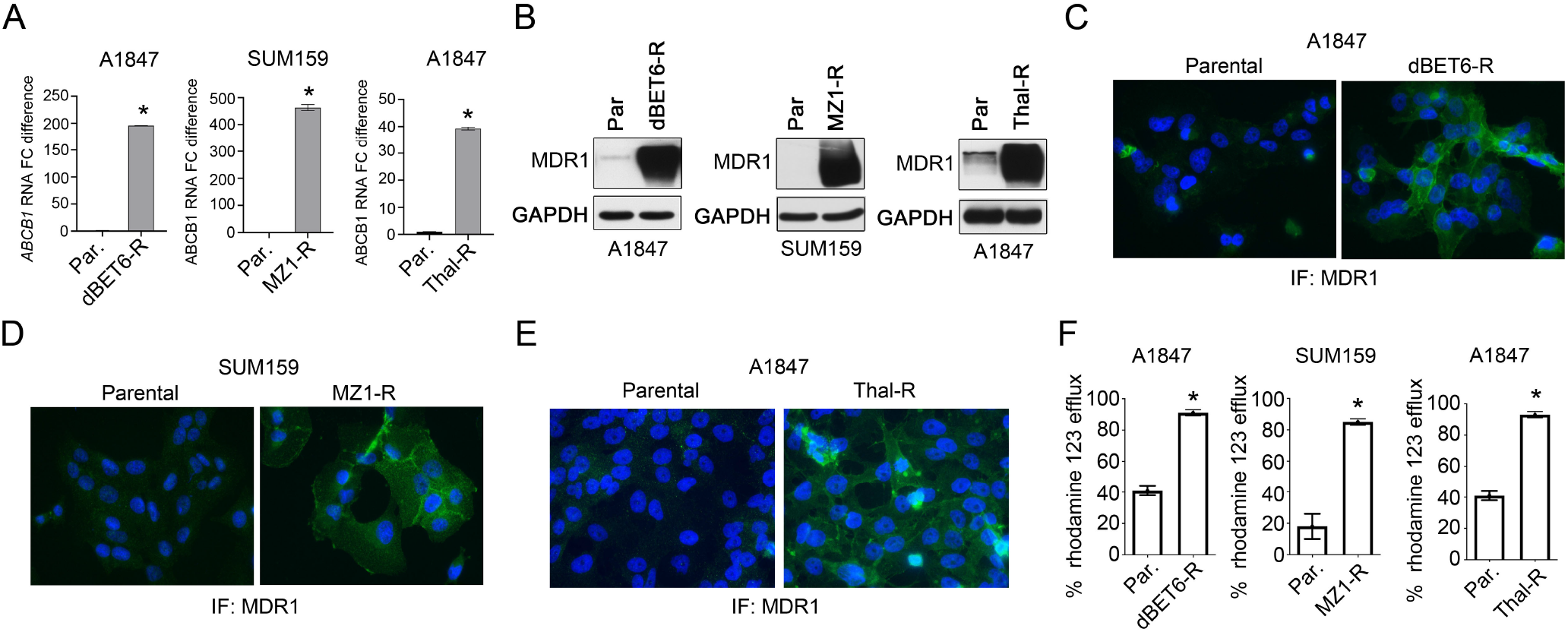
Chronic Exposure to Degraders Induces MDR1 Expression and Drug Efflux Activity. (A) ABCB1 mRNA levels are upregulated in degrader-resistant cell lines as determined by qRT-PCR. (B) MDR1 protein levels are upregulated in degrader-resistant cell lines relative to parental cells as determined by immunoblot. Blots are representative of 3 independent blots. (C-E) Confocal fluorescence microscopy of MDR1 protein levels in dBET6-R (C), MZ1-R (D) and Thal-R (E) relative to parental cell lines. MDR1 was detected by immunofluorescence using anti-MDR1 antibodies and nuclear staining by DAPI. Images are representative of 3 independent experiments. (F) Bar graph depicts increased drug efflux activity in dBET6-R, MZ1-R and Thal-R cells relative to parental cells. MDR1 drug efflux activity was measured using Rhodamine 123 efflux assays. Data present in (A), (F), are triplicate experiments SD. *p ≤0.05 by student’s t-test. Also see Figure S2.

### Genetic depletion or small molecule inhibition of MDR1 re-sensitizes degrader-resistant cells to PROTACs

Elevated levels of MDR1 have been shown to promote drug resistance in cancer cells via efflux of large hydrophobic molecules, such as chemotherapy agents (*22*). Notably, BET protein or CDK9 degrader-resistant cells acquired resistance to paclitaxel (**Fig. S3A**), a known substrate of MDR1 (*22*), as well as were cross-resistant to PROTACs targeting other proteins (**Fig. S3B-C**). Knockdown of *ABCB1* reduced cell viability in dBET6-R or Thal-R A1847 cells (**Fig. 3A**) or MZ1-R SUM159 cells (**Fig. 3B**) while exhibiting minimal effects in parental cells, demonstrating degrader-resistant cells acquired dependency on MDR1 for survival. Moreover, genetic depletion of *ABCB1* restored degradation of BET proteins or CDK9 in degrader-resistant cells, re-sensitizing cells to the degraders causing apoptosis (**Fig. 3C-E**). In contrast, knockdown of *ABCB1* in parental cells showed no effect on BET proteins or CDK9 protein levels nor induced PARP cleavage that was observed in degrader-resistant cells.

**Figure 3.**
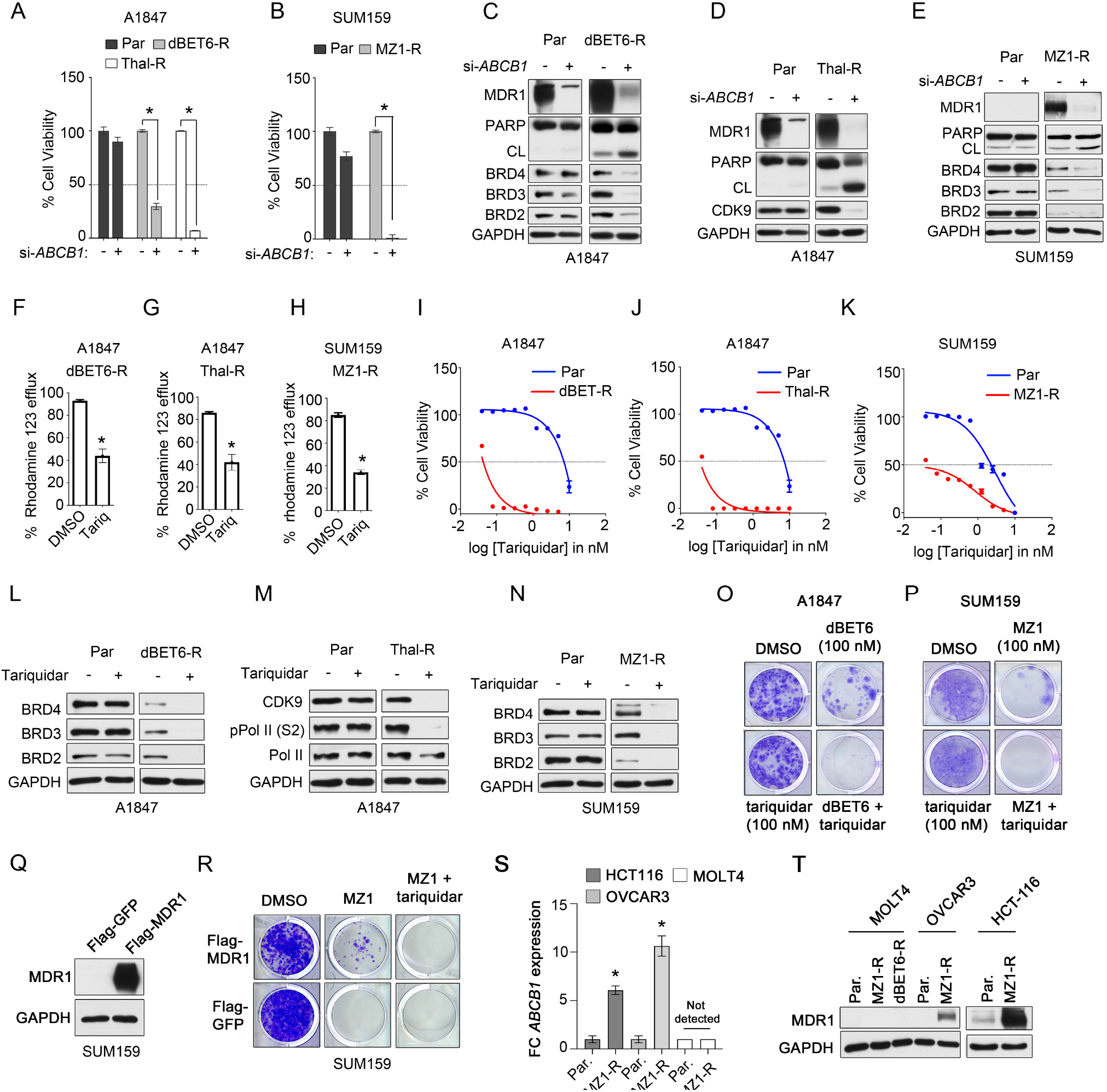
Blockade of MDR1 Activity Re-Sensitizes Degrader-Resistant Cells to PROTACs. (A-B) Degrader-resistant cells acquire dependency on MDR1 for survival. Cell-Titer Glo assay for cell viability of parental, dBET6-R or Thal-R A1847 cells (A) or parental or MZ1-R SUM159 cells (B) transfected with siRNAs targeting ABCB1 or with control siRNA and cultured for 120 hours. (C-E) Knockdown of ABCB1 in dBET6-R (C) or Thal-R (D) A1847 cells or in MZ1-R SUM159 cells (E) promotes degradation of PROTAC-targets. A1847 parental, dBET6-R or Thal-R cells were transfected with siRNAs targeting ABCB1 or with control siRNA and proteins measured by western blot. Blots are representative of 3 independent blots. (F-H) Treatment of degrader-resistant cells with tariquidar reduces MDR1 activity. Bar graph depicts decreased drug efflux activity in dBET6-R (F) or Thal-R (G) A1847 cells or MZ1-R SUM159 cells (H) relative to parental cells. Cells were treated with 0.1 μM tariquidar and MDR1 drug efflux activity was measured using Rhodamine 123 efflux assays. (I-K) Degrader-resistant cells exhibit increased sensitivity to MDR1 inhibitors. Cell-Titer Glo assay for cell viability of parental, dBET6-R (I) or Thal-R (J) A1847 cells or parental or MZ1-R SUM159 cells (K) with increasing concentrations of MDR1 inhibitor tariquidar. (L-N) Treatment of parental, dBET6-R (L) or Thal-R (M) A1847 cells or parental or MZ1-R SUM159 cells (N) promotes degradation of PROTAC-targets. A1847 parental, dBET6-R or Thal-R cells or SUM159 parental or MZ1-R cells were treated with tariquidar (0.1 μM) for 24 hours and proteins measured by western blot. Blots are representative of 3 independent blots. (O-P) MDR1 inhibition blocks development of degrader-resistance. A1847 cells were treated with DMSO, tariquidar (0.1 μM), dBET6 (0.1 μM) or the combination and colony formation assessed following 14-days of treatment (O). SUM159 cells were treated with DMSO, tariquidar (0.1 μM), MZ1 (0.1 μM) or the combination and colony formation assessed following 14-days of treatment (P). Colony formation image representative of 3 independent assays. (Q) Forced expression of Flag-MDR1 in SUM159 cells. SUM159 cells were transfected with Flag-MDR1 and selected with hygromycin. MDR1 protein expression was verified by western blot. (R) Forced expression of Flag-MDR1 promotes resistance to dBET6. SUM159 cells expressing Flag-MDR1 were treated with DMSO, MZ1 (0.1 μM), or MZ1 (0.1 μM) and tariquidar (0.1 μM) and colony formation assessed following 14 days of treatment by crystal violet staining. Colony formation image representative of 3 independent assays. (S-T) MOLT4 cells do not induce *ABCB1* expression following chronic exposure to MZ1 that is observed with OVCAR3 and HCT116. ABCB1 expression and protein levels were assessed in parental or MZ1-R cells using qRT-PCR (S) or immunoblot (T). Blots are representative of 3 independent blots. Data present in (A), (B), (F-H), (I-K), and (S) are triplicate experiments SD. *p ≤0.05 by student’s t-test. Also see Figure S3.

Several small molecule inhibitors of MDR1 have been developed, including tariquidar (*23*), which is currently being evaluated in clinical trials for the treatment of MDR1-driven drug resistant disease (*24*). Treatment of A1847 dBET6-R, Thal-R or MZ1-R SUM159 cells with tariquidar reduced MDR1 drug efflux pump activity, indicated by reduced efflux of Rhodamine 123 in degrader-resistant cells compared to parental cells (**Fig. 3F-H**). Moreover, degrader-resistant cells were more sensitive to tariquidar than parental cells (**Fig. 3I-K**), and inhibition of MDR1 function restored degradation of BET proteins or CDK9 (**Fig. 3L-N**). Notably, chronic exposure of A1847 cells to BET inhibitor JQ1 did not cause sensitization to tariquidar, suggesting that acquired dependency on MDR1 was distinct to degrader-resistance (**Fig. S3D**). Combined treatment of A1847 or SUM159 cells with BET protein degraders and tariquidar blocked the development of BET protein degrader resistant colonies over a 14-day period (**Fig. 3O-P**). Moreover, forced expression of Flag-MDR1 in SUM159 cells rescued colony formation growth in MZ1-treated cells that could be blocked by tariquidar treatment, signifying overexpression of MDR1 reduces sensitivity towards BET degraders (**Fig. 3Q-R**).

To further explore MDR1 upregulation in degrader-resistance in cancer cells, we chronically exposed 3 additional cancer cell lines (OVCAR3, HCT116 and MOLT4) to BET protein degraders and assessed MDR1 protein levels. OVCAR3 and HCT116 cell lines acquired resistance to MZ1 (**Fig. S3E-F**) that was accompanied by elevated MDR1 mRNA and protein levels in parental cells (**Fig. 3S-T**), as well as an increased sensitivity towards tariquidar-treatments (**Fig.S3G-H**). In contrast, we were unable to generate MZ1-resistant MOLT4 cells (**Fig. 3SI**) and chronic exposure to BET protein degraders did not result in upregulation of *ABCB1* mRNA or protein levels (**Fig. 3S-T**). These findings suggest that not all cancer cells will induce MDR1 following continuous degrader exposure, in our studies, 4 out of 5 cancer cell lines induced MDR1.

Together, our findings demonstrate cancer cells can acquire resistance to degrader therapies through upregulation of the multidrug resistance pump MDR1 and inhibition of MDR1 restores degrader function overcoming drug resistance in degrader-resistant cancer cells.

### MDR1 overexpressing cells exhibit intrinsic resistance to PROTAC therapies that can be overcome by MDR1 inhibition

Overexpression of MDR1 frequently occurs in cancers conveying intrinsic resistance to several anti-cancer therapies such as chemotherapies (*19*). Analysis of *ABCB1* mRNA expression across the cancer cell line encyclopedia (*25, 26*) revealed colorectal, neuroblastoma, hepatobiliary and renal cell carcinomas exhibited frequent overexpression of MDR1 (**Fig. S4A**). Moreover, querying the human protein atlas, elevated MDR1 protein levels were observed in >50% of liver and colorectal cancer tumors by immunohistochemistry (IHC) (*27*) (**Fig. S4B**). To determine if overexpression of MDR1 in cancer cell lines influences degrader-sensitivity, we queried a prior study which explored MZ1 or dBET6 resistance across a panel of various cancer cell lines (*11*) with publicly available *ABCB1* mRNA expression datasets (*28*). Notably, cancer cell lines that were resistant to both MZ1 and dBET6 expressed *ABCB1* at higher levels than those sensitive to the degraders (P<0.001), suggesting *ABCB1* expression represents a potential biomarker for BET protein degrader response in cancer cells (**Fig. 4A**).

**Figure 4.**
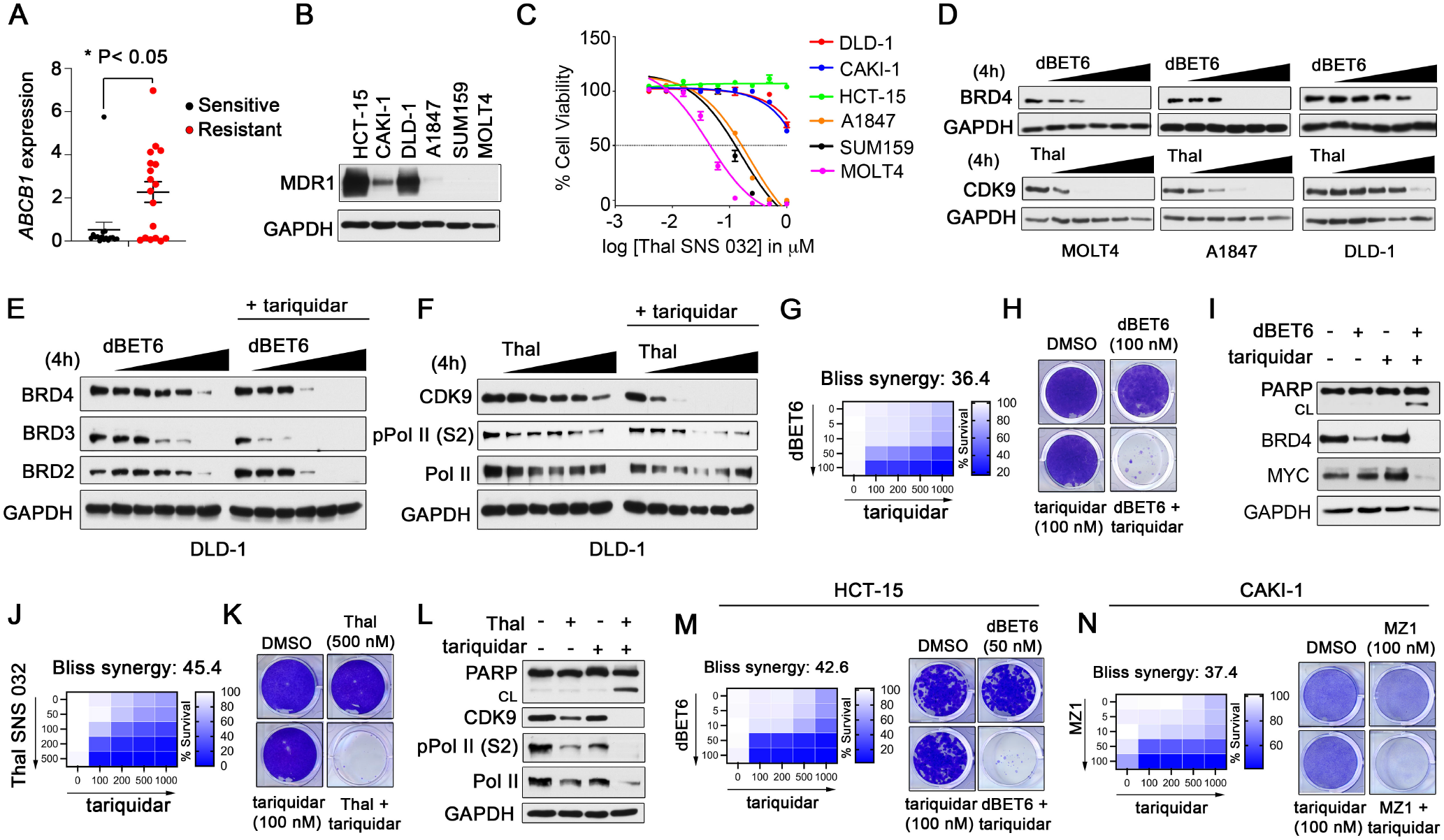
Overexpression of MDR1 Conveys Intrinsic Resistance to Degrader Therapies in Cancer Cells. (A) Cancer cells resistant to BET protein degraders harbor elevated *ABCB1* expression. Expression of *ABCB1* in cancer cell lines exhibiting sensitivity or resistance to MZ1/dBET6 was queried from (*25*) and sensitivity or resistance to degraders obtained from (*9*). Difference in ABCB1 expression amongst degrader-resistant or sensitive was determined by students t-test. (B) MDR1 protein levels in a panel of cancer cell lines as determined by western blot. Blots are representative of 3 independent blots. (C) Cancer cells overexpressing MDR1 exhibit reduced sensitivity towards Thal SNS 032. Cancer cells were treated with escalating doses of Thal SNS 032 for 5 d and cell viability assessed by CellTiter-Glo. (D) Overexpression of MDR1 reduces PROTAC-mediated degradation efficiency in cancer cells. Cancer cells exhibiting different levels of MDR1 were treated with escalating doses of dBET6 or Thal SNS 032 (Thal) for 4 hours and BRD4 or CDK9 protein levels assessed by western blot. Blots are representative of 3 independent blots. (E-F) Combined inhibition of MDR1 improves PROTAC-mediated degradation in MDR1 overexpressing cells. DLD-1 cells were treated with increasing doses of dBET6 alone or in combination with tariquidar (0.1 μM) (E) or increasing doses of Thal SNS 032 alone or in combination with tariquidar (0.1 μM) (F) for 4 hours and BRD4 or CDK9 protein levels assessed by western blot. Blots are representative of 3 independent blots. (G-I) Combining tariquidar and dBET6 exhibits drug synergy in MDR1-overexpressing cells. Cell-Titer Glo assay for cell viability of DLD-1 cells treated with increasing concentrations of dBET6, tariquidar or the combination and bliss synergy scores determined (G). DLD-1 cells were treated with DMSO, tariquidar (0.1 μM), dBET6 (0.1 μM) or the combination and colony formation assessed following 14 days of treatment (H). Colony formation image representative of 3 independent assays. Western blot analysis was performed on DLD-1 cells treated with DMSO, tariquidar (0.1 μM), dBET6 (0.1 μM) or the combination for 24 hours (I). Blots are representative of 3 independent blots. (J-L) Combining tariquidar and Thal SNS 032 exhibits drug synergy in MDR1-overexpressing cells. Cell-Titer Glo assay for cell viability of DLD-1 cells treated with increasing concentrations of Thal SNS 032, tariquidar or the combination and Bliss synergy scores determined (J). DLD-1 cells were treated with DMSO, tariquidar (0.1 μM), Thal SNS 032 (0.5 μM) or the combination and colony formation assessed following 14 days of treatment (K). Colony formation image representative of 3 independent assays. Western blot analysis was performed on DLD-1 cells treated with DMSO, tariquidar (0.1 μM), Thal SNS (0.5 μM) or the combination for 24 hours (L). Blots are representative of 3 independent blots. (M-N) Combining tariquidar with BET degraders enhances growth inhibition of MDR1-overexpressing cell lines HCT-15 and CAKI-1. Cell-Titer Glo assay for cell viability of cells treated with increasing concentrations of dBET6 (M) or MZ1 (N), tariquidar or the combination and bliss synergy scores determined. Cells were treated with DMSO, tariquidar (0.1 μM), dBET6 (0.05 μM) (M), MZ1 (0.1 μM) (N) or the combination and colony formation assessed following 14-days of treatment. Colony formation image representative of 3 independent assays. Data present in (C), (G), (J), (M-N) are triplicate experiments SD. *p ≤0.05 by student’s t-test. Also see Figure S4.

To further explore MDR1 as a candidate biomarker for degrader resistance, we selected 3 cancer cell lines, HCT-15 (colon), DLD-1 (colon) and CAKI-1 (renal) with established overexpression of MDR1 and compared the impact of degrader-treatment on cell viability and protein degradation with cell lines that express low (A1847) or no detectable levels of ABCB1 (SUM159 and MOLT4) by immunoblot (**Fig. 4B**). Treatment of MDR1 overexpressing cells with Thal SNS 032, MZ1, or dBET6 did not reduce cell viability to the extent of cancer cell lines expressing low or no detectable MDR1 protein (**Fig. 4C, S4C-D**). Similarly, treatment of MDR1 overexpressing cell line DLD-1 with dBET6 or Thal SNS 032 did not reduce the intended degrader target to the extent observed with degrader-sensitive A1847 or MOLT4 cells (**Fig. 4D**). Importantly, co-treatment of DLD-1 cells with tariquidar and either dBET6 (**Fig. 4E**) or Thal SNS 032 (**Fig. 4F**) improved the degradation efficiency, resulting in a greater reduction in BET proteins or CDK9 at lower concentrations of the PROTACs. Additionally, co-treatment of DLD-1 cells with FAK degrader (FAK-degrader-1) (*29*) or MEK1/2 degrader (MS432) (*30*) and tariquidar improved the protein reduction relative to single agent therapies (**Fig. S4E-F**), suggesting overexpression of MDR1 promotes resistance to degrader therapies, independent of protein target.

Combination therapies involving BET protein degraders and tariquidar in DLD-1 cells exhibited high drug synergy (Bliss synergy score 36.4) in blocking cell viability in 5-day growth assays and inhibited colony formation over a 14-day period better than single agent therapies (**Fig. 4G-H**). Moreover, co-administration of dBET6 and tariquidar improved protein degradation of BET proteins, reduced the expression of the BRD4 target MYC and induced apoptosis (**Fig. 4I**). Similarly, co-treatment of DLD-1 cells with tariquidar and Thal SNS 032 blocked cell viability, and colony formation to a greater extent than single agent therapies, as well as reduced CDK9 and CDK9-substrate Pol II (S2) and induced apoptosis uniquely in the combination therapy (**Fig. 4J-L**). The drug synergy amongst tariquidar and BET protein or CDK9 degraders was also observed in additional MDR1 overexpressing cell lines HCT-15 (**Fig. 4M, S4G**) and CAKI-1 (**Fig. 4N, S4H**). Together, our findings suggest specific types of cancers that express high levels of MDR1 such as colorectal or renal cancers will likely exhibit intrinsic resistance to degraders requiring co-administration of MDR1 inhibitors to achieve protein degradation and therapeutic efficacy.

### Repurposing dual kinase/MDR1 inhibitors to overcome degrader-resistance in cancer cells

Specific inhibitors of MDR1 such as tariquidar have shown limited success in the clinic at re-sensitizing MDR1 overexpressing patients to chemotherapy due to toxicities, low drug-drug interactions and the inability to achieve desired concentrations of tariquidar in tumors (*31*). Notably, several kinase inhibitors have been shown to be potent inhibitors of MDR1 drug efflux activity capable of overcoming multidrug resistance in cancer cells (*32*). The ErbB receptor inhibitor lapatinib is an FDA approved drug for the treatment of several HER2 driven cancers and has been shown to also directly inhibit MDR1 drug efflux activity both in cancer cells and *in vivo* tumor models (*33*). Additionally, RAD001, an FDA approved mTORC1 inhibitor for treatment of renal cell carcinomas, has also been shown to inhibit MDR1 function in cancer cells (*34*). Based on these findings, we hypothesized that the combined inhibition of ErbB receptors or mTORC1 and MDR1 drug efflux by lapatinib or RAD001 could represent a promising strategy to overcome MDR1-mediated resistance to degraders, as well as improve anti-cancer benefits of PROTACs.

Treatment of degrader-resistant cell lines dBET6-R or Thal-R cell lines with RAD001 or lapatinib reduced MDR1 drug efflux activity similar to that observed with tariquidar (**Fig. 5A-B**). Degrader-resistant cell lines were more sensitive to RAD001 (**Fig. 5C-D, S5A-B**) or lapatinib (**Fig. 5E-F, S5C-D**) than parental cells and administration of RAD001 or lapatinib resulted in degradation of BET proteins (**Fig. 5G, S5E-F**) or CDK9 (**Fig. 5H**) uniquely in degrader-resistant cell lines. Moreover, treatment of BET protein (**Fig. 5I**) or CDK9 (**Fig. 5J**) degrader-resistant cell lines with RAD001 or lapatinib resulted in apoptosis similar to tariquidar treatment, demonstrating RAD001 or lapatinib can block MDR1 function overcoming MDR1-driven degrader-resistance.

**Figure 5.**
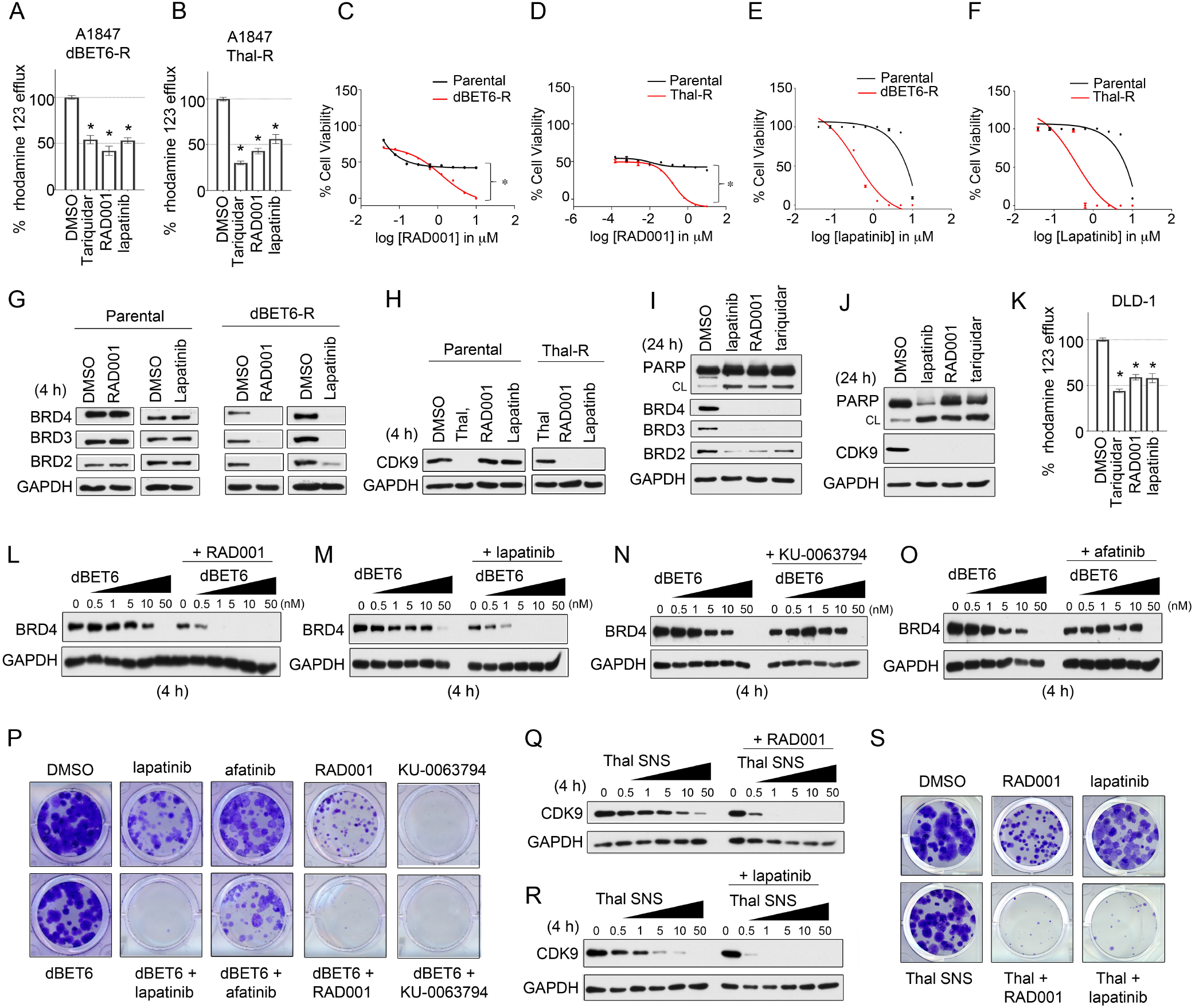
Re-Purposing Dual Kinase/MDR1 Inhibitors to Overcome Degrader Resistance in Cancer Cells. (A-B) Treatment of degrader-resistant cells with RAD001 or lapatinib reduces MDR1 drug efflux activity. A1847 parental, dBET6-R (A) or Thal-R (B) cells were treated with DMSO, 2 μM tariquidar, 2 μM RAD001, or 2 μM lapatinib and Rhodamine 123 efflux assessed. (C-D) Degrader-resistant cells exhibit increased sensitivity towards RAD001. Cell-Titer Glo assay for cell viability of A1847 parental, dBET6-R (C) or Thal-R (D) cells treated with increasing concentrations of RAD001. (E-F) Degrader-resistant cells exhibit increased sensitivity towards lapatinib. Cell-Titer Glo assay for cell viability of A1847 parental, dBET6-R (C) or Thal-R (D) cells treated with increasing concentrations of lapatinib. (G-H) Treatment of degrader-resistant cells with RAD001 or lapatinib promotes degradation of PROTAC-targets. A1847 parental, dBET6-R (G) or Thal-R (H) cells treated with DMSO, RAD001 (2 μM) or lapatinib (2 μM) for 4 hours and proteins measured by western blot. Blots are representative of 3 independent blots. (I-J) Treatment of degrader-resistant cells with RAD001 or lapatinib induces apoptosis. A1847 parental, dBET6-R (I) or Thal-R (J) cells treated with DMSO, RAD001 (2 μM), lapatinib (2 μM) or tariquidar (2 μM) for 24 hours and proteins measured by western blot. Blots are representative of 3 independent blots. (K) Treatment of MDR1-overexpressing cells with RAD001 or lapatinib reduces MDR1 drug efflux. DLD-1 cells were treated with DMSO, 2 μM tariquidar, 2 μM RAD001, or 2 μM lapatinib and Rhodamine 123 efflux assessed. (L-M) Combined RAD001 or lapatinib-treatment improves PROTAC-mediated degradation of BRD4 in MDR1 overexpressing cells. DLD-1 cells were treated with increasing doses of dBET6 alone or in combination with RAD001 (2 μM) (L) or lapatinib (2 μM) (M) for 4 hours and BRD4 protein levels assessed by western blot. Blots are representative of 3 independent blots. (N-O) KU-0063794 or Afatinib do not improve PROTAC-mediated degradation of BRD4 in MDR1 overexpressing cells. DLD-1 cells were treated with increasing doses of dBET6 alone or in combination with KU-0063794 (2 μM) (N) or afatinib (2 μM) (O) for 4 hours and BRD4 protein levels assessed by western blot. Blots are representative of 3 independent blots. (P) Combining RAD001 or lapatinib but not KU-0063794 or Afatinib with BET degraders exhibits drug synergy in MDR1-overexpressing cells. DLD-1 cells were treated with DMSO, dBET6 (0.1 μM), lapatinib (2 μM), afatinib (2 μM), RAD001 (2 μM), KU-0063794 (2 μM) or in combination with dBET6 and colony formation assessed following 14 days of treatment. Colony formation image representative of 3 independent assays. (Q-R) Combined RAD001 or lapatinib-treatment improves PROTAC-mediated degradation of CDK9 in MDR1 overexpressing cells. DLD-1 cells were treated with increasing doses of Thal SNS 032 alone or in combination with RAD001 (2 μM) (L) or lapatinib (2 μM) (M) for 4 hours and CDK9 protein levels assessed by western blot. Blots are representative of 3 independent blots. (S) Combining RAD001 or lapatinib with CDK9 degraders exhibits drug synergy in MDR1-overexpressing cells. DLD-1 cells were treated with DMSO, dBET6 (0.1 μM), lapatinib (2 μM), RAD001 (2 μM) or in combination with Thal SNS 032 and colony formation assessed following 14 days of treatment. Colony formation image representative of 3 independent assays. Data present in (C-F), and (K) are triplicate experiments SD. *p ≤0.05 by student’s t-test. Also see Figure S5.

Next, we explored whether RAD001 or lapatinib treatment could sensitize MDR1-overexpressing cells to degrader therapies. Treatment of DLD-1 cells with RAD001 or lapatinib reduced MDR1 drug efflux activity similar to tariquidar treatment (**Fig. 5K**), and immunoblot analysis showed RAD001 or lapatinib treatment improved dBET6-mediated degradation of BRD4 lowering the concentration of dBET6 required to achieve maximal protein degradation (**Fig. 5L-M**). Notably, a 100-fold reduction in concentrations of dBET6 were required to degrade BRD4 when combined with RAD001 or lapatinib. In contrast, combined treatment of DLD-1 cells with KU-0063794 (MTOR inhibitor) or afatinib (ErbB receptor inhibitor), drugs that do not inhibit MDR1 function (**Fig. S5G**), failed to improve degradation of BRD4 (**Fig. 5N-O**). Moreover, treatment of DLD-1 cells with lapatinib but not afatinib sensitized DLD-1 cells to dBET6 providing durable inhibition of colony formation over a 14-day period (**Fig. 5P**). Similarly, co-treatment of DLD-1 cells with KU-0063694 and dBET6 did not improve growth inhibition observed with the RAD001 and dBET6 combination, where single agent KU-0063694 treatment completely repressed colony formation. RAD001 or lapatinib treatment also sensitized DLD-1 cells to Thal SNS 032, improving degradation of CDK9 (**Fig. 5Q-R**), and enhancing growth inhibition of colonies (**Fig. 5S**). Together, these findings demonstrate RAD001 or lapatinib can be utilized as MDR1 inhibitors to overcome degrader-resistance mediated by MDR1 drug efflux.

### Lapatinib-treatment enhances MEK1/2 degrader therapies in K-ras mutant colorectal cancer cells by dual blockade of MDR1 activity and ERBB receptor signaling

K-ras mutations occur in nearly 40% of colorectal cancer (CRC) patients, supporting therapies that target K-ras effectors such as the MEK-ERK signaling pathway (*35*). Recently, MEK1/2 degraders have been developed that show potent anti-growth properties in RAS-RAF altered cancers (*30*). Notably, the majority of K-ras mutant CRC cell lines exhibit elevated *ABCB1* expression (*28*), suggesting concomitant blockade of MDR1 may be required to achieve therapeutic efficacy with MEK1/2 degraders (**Fig. 6A-B**). Moreover, resistance to MEK inhibitors in K-ras mutant colorectal cancer cells is mediated by activation of ErbB receptors and downstream RAF-MEK-ERK and PI3K/AKT signaling (*36*). Based on these findings, we hypothesized combination therapies involving lapatinib and MEK1/2 degrader MS432 could be a unique strategy to simultaneously block MDR1-mediated resistance, as well as inhibit MEKi-mediated kinome reprogramming involving activation of ERBB3 signaling.

**Figure 6.**
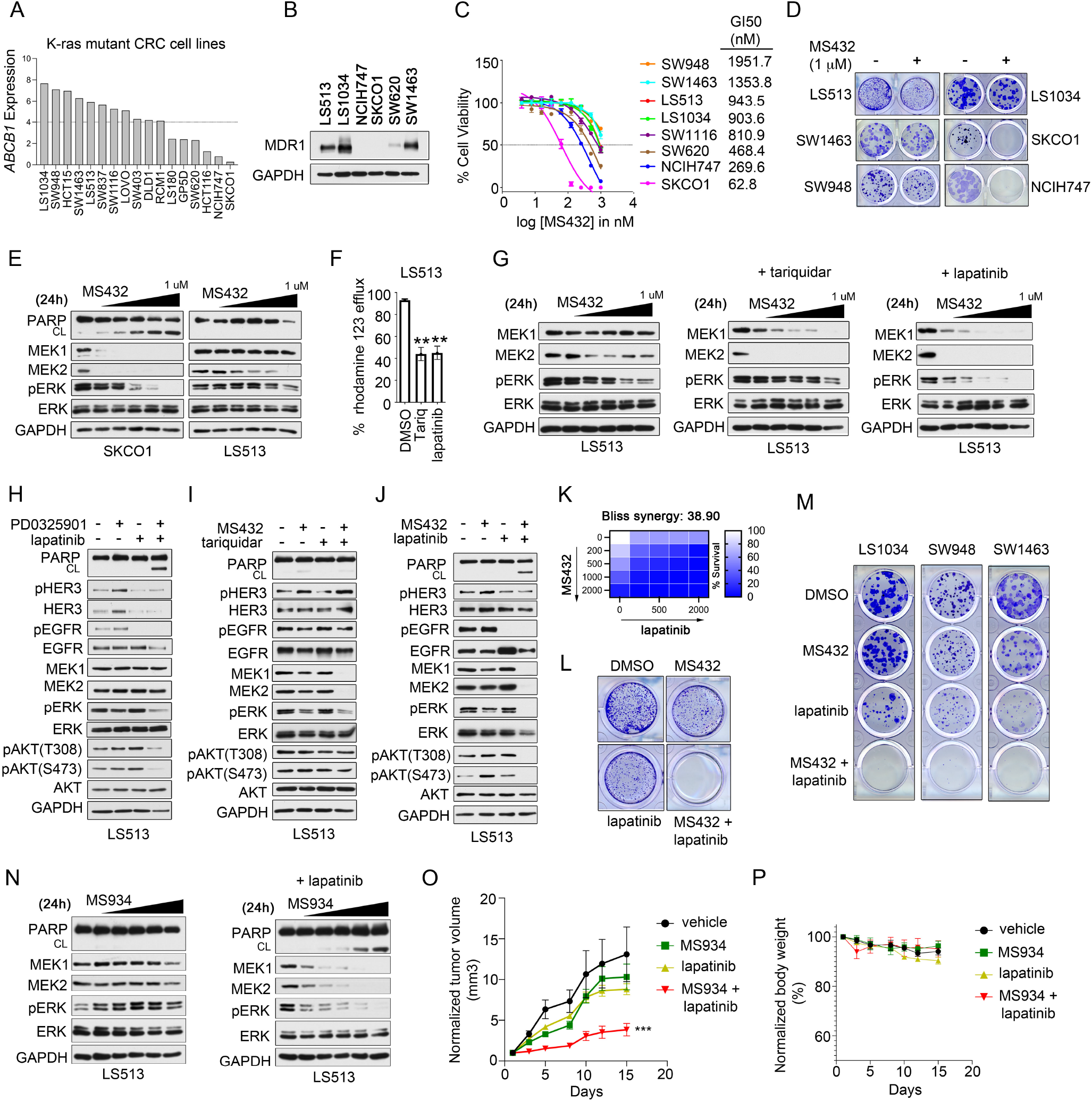
Combining MEK1/2 Degraders with Lapatinib Synergize to Kill MDR1-Overexpressing K-ras Mutant CRC Cells and Tumors. (A-B) MDR1 is overexpressed in the majority of K-ras mutant CRC cell lines. (A) *ABCB1* expression data was obtained from c-Bioportal. (B) MDR1 protein levels across selected CRC cell lines was determined by western blot. Blots are representative of 3 independent blots. (C-D) K-ras mutant CRC cells overexpressing MDR1 exhibit reduced sensitivity towards MEK1/2 degrader MS432. (C) CRC cells were treated with escalating doses of MS432 for 5 d and cell viability assessed by CellTiter-Glo. GI50 values were determined in Prism software. (D) CRC cells were treated with 1 μM of MS432 and colony formation assessed following 14 days of treatment. Colony formation image representative of 3 independent assays. (E) Overexpression of MDR1 reduces PROTAC-mediated degradation efficiency in K-ras mutant CRC cells. CRC cells exhibiting different levels of MDR1 were treated with escalating doses of MS432 for 4 hours and MEK1/2 protein levels assessed by western blot. Blots are representative of 3 independent blots. (F) Treatment of MDR1-overexpressing cells with tariquidar or lapatinib reduces MDR1 drug efflux. DLD-1 cells were treated with DMSO, 2 μM tariquidar, or 2 μM lapatinib and Rhodamine 123 efflux assessed. (G) Combined inhibition of MDR1 improves PROTAC-mediated degradation in MDR1 overexpressing cells. LS513 cells were treated with increasing doses of MS432 alone or in combination with tariquidar (0.1 μM) or increasing doses of MS432 alone or in combination with lapatinib (5 μM) for 24 hours and protein/phosphoprotein levels assessed by western blot. Blots are representative of 3 independent blots. (H) MEK inhibition upregulates ErbB receptor signaling and downstream AKT signaling in LS513 cells that can be blocked by lapatinib. LS513 cells were treated with DMSO, PD0325901 (0.01 μM), lapatinib (5 μM), or the combination for 48 hours and signaling assessed by western blot. Blots are representative of 3 independent blots. (I-J) Lapatinib but not tariquidar treatment blocks MEKi-induced ERBB3 reprogramming. LS513 cells were treated with DMSO, MS432 (1 μM), tariquidar (0.1 μM) or the combination (I) or DMSO, MS432 (1 μM), lapatinib (5 μM) or the combination (J) and protein/phosphoproteins assessed by western blot. Blots are representative of 3 independent blots. (K-L) Combining lapatinib and MS432 exhibits drug synergy in MDR1-overexpressing K-ras mutant CRC cells. Cell-Titer Glo assay for cell viability of LS513 cells treated with increasing concentrations of MS432, lapatinib or the combination of lapatinib and MS432 (H). Bliss synergy scores determined. LS513 cells were treated with DMSO, lapatinib (2 μM), MS432 (1 μM) or the combination and colony formation assessed following 14 days of treatment (I). Colony formation image representative of 3 independent assays. (M) Lapatinib in combination with MS432 enhances growth inhibition in MDR1-overexpressing K-ras mutant CRC cell lines. CRC cell lines were treated with DMSO, lapatinib (2 μM), MS432 (1 μM), or the combination and colony formation assessed following 14 days of treatment. Colony formation image representative of 3 independent assays. (N) Co-treatment with MS934 and lapatinib MDR1 improves PROTAC-mediated degradation in MDR1 overexpressing cells. LS513 cells were treated with increasing doses of MS934 alone or in combination with lapatinib (5 μM) for 24 hours and protein/phosphoprotein levels assessed by western blot. Blots are representative of 3 independent blots. (O-P) MEK degraders in combination with lapatinib reduce tumor growth *in vivo*. LS513 cells were grown as xenografts in nude mice and treated with vehicle, 50 mg/kg MS934, 100 mg/kg lapatinib, or the combination of MS934 and lapatinib and tumor volume determined (O). Body weight of animals was determined to evaluate potential toxicities of drug treatments (P). N=5 per treatment group, Error bar ± SEM. Data present in (C), (F), (K) and (L) are triplicate experiments SD. *p ≤0.05 by student’s t-test. Also see Figure S6.

As predicted, MDR1 overexpressing K-ras mutant CRC cell lines (LS1034, LS513, SW948 and SW1463) were more resistant to MEK1/2 degrader MS432 than MDR1 low expressing CRC cell lines (SKCO1, NCIH747, and SW620) (**Fig. 6C-D**). Notably, all K-ras mutant cell lines were sensitive to treatment with MEK inhibitor, trametinib (*37*) (**Fig. S6A**). Moreover, treatment of MDR1-overexpressing cell line LS513 with MS432 did not reduce MEK1 or MEK2 protein levels, inhibit ERK1/2 phosphorylation or induce apoptosis that was observed with degrader-sensitive MDR1 non-expressing cell line SKCO1 (**Fig. 6E**). Treatment of LS513 cells with lapatinib reduced MDR1 drug efflux activity similar to tariquidar (**Fig. 6F**), and co-treatment of LS513 cells with MS432 and lapatinib improved the degradation efficiency of MEK1 and MEK2, as well as reduced ERK1/2 phosphorylation at lower concentrations of MS432 (**Fig. 6G**). Notably, the addition of lapatinib to MS432 reduced levels of ERK1/2 activating phosphorylation to a greater extent than the tariquidar/MS432 combined treatment, suggesting concurrent blockade of ErbB receptors and MDR1 may be more efficacious than inhibiting MDR1 activity alone.

Next, we explored the impact of blockade of MDR1 alone using tariquidar or MDR1 and ErbB receptors using lapatinib on K-ras effector signaling in LS513 cells. As previously reported, treatment of LS513 cells with MEK inhibitors induced ERBB3 and downstream AKT and RAF signaling, which could be blocked by lapatinib treatment (**Fig. 6H**), and combining lapatinib and PD0325901 exhibited drug synergy (**Fig. S6B**). Notably, co-treatment of LS513 cells with MS432 and lapatinib but not tariquidar reduced MEKi-induced ERBB3 and downstream AKT activation, as well as distinctly induced apoptosis (**Fig. 6I-J**). Combination therapies involving MS432 and lapatinib in LS513 cells exhibited robust drug synergy with a Bliss synergy score of 38.9 (**Fig. 6K)**, as well as provided durable inhibition of colony formation over a 14-day period (**Fig. 6L**). Furthermore, the combination of lapatinib and MS432 provided durable growth inhibition of other MDR1-overexpressing K-ras mutant CRC cell lines (**Fig. 6M**). Next, we explored the efficacy of combining MEK degraders and lapatinib *in vivo* using LS513 xenograft models and the recently published MEK degrader MS934, which has optimal bioavailability for animal studies (*30*). Similar to MS432, combining MS934 and lapatinib enhanced MEK1/2 degradation in LS513 cells, exhibited drug synergy, and distinctly induced apoptosis (**Fig. 6N, S6C**). Treatment of mice harboring LS513 xenografts with the MEK degrader MS934 and lapatinib distinctly reduced tumor growth with minimal impact on mice body weight, while single agents were ineffective (**Fig. 6O-P**), suggesting concurrent blockade of ErbB receptors and MDR1 will likely be required to achieve therapeutic response using MEK degraders in K-ras mutant CRC.

### Combining lapatinib and KRAS^G12C^ degrader LC-2 exhibits drug synergy in K-ras G12C mutant CRC cells

PROTACs targeting KRAS^G12C^ mutants have recently been developed that induce rapid and sustained degradation of KRAS^G12C^ leading to inhibition of MAPK signaling in KRAS^G12C^ cancer cell lines (*7*). Notably, several KRAS^G12C^ cancer cell lines have been shown to be resistant to KRAS^G12C^ inhibitors but sensitive to K-ras knockdown (*38*), suggesting degradation of KRAS^G12C^ may be an alternative therapeutic strategy for these K-ras inhibitor-resistant cells. However, similar to MEK1/2 inhibitors, adaptive resistance to KRAS^G12C^ inhibitors in CRC cells has also been shown to mediated by kinome remodeling involving activation of ErbB receptor signaling bypassing K-ras inhibition (*39*). Here, we explored whether combining lapatinib and the KRAS^G12C^ degrader LC-2 (*7*), would improve degradation efficiency of KRAS^G12C^ and enhance therapeutic efficacy in MDR1-overexpressing KRAS^G12C^ CRC cell lines, SW1463 (homozygous KRAS^G12C^) and SW837 (heterozygous KRAS^G12C^).

SW1463 or SW837 KRAS^G12C^ CRC cells exhibited intrinsic resistance to LC-2 but were sensitive to KRAS^G12C^ inhibitor MRTX849 treatment (**Fig. 7A-B**). Treatment of SW1463 cells with 1 μM LC-2 had no impact on KRAS^G12C^ protein levels, while combining tariquidar or lapatinib with LC-2 improved PROTAC-mediated degradation of KRAS^G12C^ reducing protein levels (**Fig. 7C-D**). Of particular interest, combining either tariquidar or lapatinib with LC-2 reduced phosphorylation of MEK and ERK, but the lapatinib combination uniquely reduced CRAF and AKT phosphorylation, as well as induced apoptosis. Similarly, co-treatment of SW837 cells with LC-2 and lapatinib but not single agents reduced KRAS effectors CRAF, AKT, MEK and ERK phosphorylation, as well as caused apoptosis (**Fig. 7E**). Notably, it was difficult to observe enhanced reduction in KRAS^G12C^ protein levels in response to LC-2 and lapatinib treatment in SW837 cells, likely due to SW837 cells expressing KRAS^WT^, which is not targeted by LC-2. Combining LC-2 and lapatinib exhibited drug synergy in SW1463 and SW837 with Bliss synergy scores of 26.8 and 25.0 (**Fig. 7F-G**), while tariquidar showed marginal synergy in either cell line (**Fig. S7A-B**). Furthermore, LC-2 in combination with lapatinib blocked colony formation in SW1463 and SW837 cells to a greater extent than LC-2/tariquidar treatments (**Fig. 7H-I**), demonstrating combined blockade of ErbB receptors and MDR1 was required to achieve durable growth inhibition using LC-2 in MDR1-overexpressing KRAS^G12C^ CRC cells.

**Figure 7.**
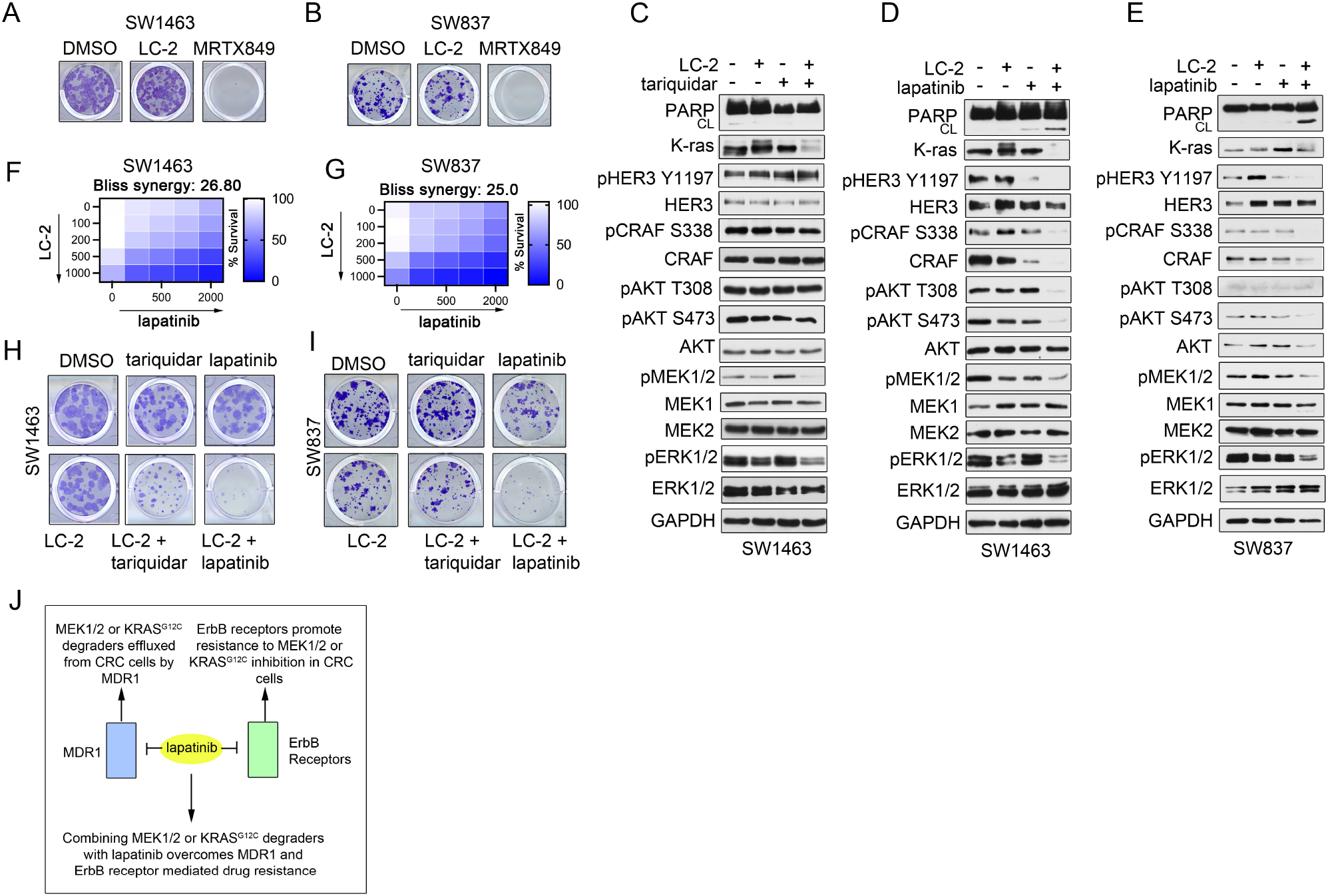
Lapatinib-treatment improves KRAS^G12C^ degrader therapies in MDR1-overexpressing CRC cell lines. (A-B) MDR1-overexpressing KRAS^G12C^ mutant CRC cell lines are resistant to LC-2 but sensitive to K-ras inhibitors. SW1463 or SW837 cell lines were treated with DMSO, LC-2 (1 μM) or MRTX849 (1 μM) and colony formation assessed following 14 days of treatment. Colony formation image representative of 3 independent assays. (C-D) Lapatinib in combination with LC-2 but not tariquidar inhibits KRAS^G12C^ effector signaling. SW1463 cells were treated with DMSO, MS432 (1 μM), lapatinib (5 μM), tariquidar (0.1 μM) or the combination of MS432/lapatinib or MS432/tariquidar for 48 hours and protein/phosphoprotein levels assessed by western blot. Blots are representative of 3 independent blots. (E) Combination therapies involving LC-2 and lapatinib block KRAS^G12C^ effector signaling. SW837 cells were treated with DMSO, MS432 (1 μM), lapatinib (5 μM) or the combination of MS432/lapatinib for 48 hours and protein/phosphoprotein levels assessed by western blot. (F-G) Combining lapatinib and LC-2 exhibits drug synergy in MDR1-overexpressing KRAS^G12C^ CRC cells. Cell-Titer Glo assay for cell viability of SW1463 (G) or SW837 (H) cells treated with increasing concentrations of LC-2, lapatinib or the combination and bliss synergy scores determined. (H-I) Combining lapatinib with LC-2 exhibits durable growth inhibition in MDR1-overexpressing KRAS^G12C^ CRC cells. SW1463 (I) or SW837 (J) cells were treated with DMSO, LC-2 (1 μM), lapatinib (2 μM), tariquidar (0.1 μM) or the combination of MS432/lapatinib or MS432/tariquidar and colony formation assessed following 14 days of treatment. Colony formation image representative of 3 independent assays. (J) Rationale for combining lapatinib with MEK1/2 or KRAS^G12C^ degraders in MDR1-overexpressing CRC cell lines. Simultaneous blockade of MDR1 and ErbB receptor signaling overcomes degrader resistance as well as ErbB receptor kinome reprogramming resulting in sustained inhibition of Kras effector signaling. Data present in (F-G), are triplicate experiments SD. *p ≤0.05 by student’s t-test. Also see Figure S7.

Together, our findings suggest the combination of dual MDR1/ErbB receptor inhibitor lapatinb and PROTACs targeting MEK1/2 or KRAS^G12C^ represents a promising combination therapy for MDR1-overexpressing K-ras mutant CRC cells due to simultaneous blockade of both MDR1 and ErbB receptor driven resistance programs (**Fig. 7J**).

## DISCUSSION

PROTACs have emerged as a new class of drugs for the treatment of cancer that can hijack the tumor cells own protein machinery to degrade oncogenic targets, including previously undruggable candidates (*4*). PROTACs have many advantages over traditional inhibitors and are avidly being pursued in clinical trials for several cancers (*40*). Here, using proteomics, we identified an acquired resistance mechanism to chronic PROTAC therapy that involved upregulation of the drug efflux pump MDR1. Moreover, we showed cancer cells overexpressing MDR1 exhibited intrinsic resistance to degraders. Importantly, we demonstrated blockade of MDR1 using selective or dual kinase/MDR1 inhibitors restored degrader sensitivity improving the longevity of PROTAC therapies. Notably, we discovered lapatinib may represent a promising drug to improve MEK1/2 or KRAS^G12C^ degrader efficacy in K-ras mutant CRCs due to simultaneous blockade of MDR1 and ErbB receptor mediated resistance.

Upregulation of MDR1 has been reported as the major resistance mechanism to chemotherapies such as taxols in cancer therapies (*22*). Our findings suggest MDR1 expression could represent a potential biomarker for efficacy of PROTACs in the treatment of cancer. Notably, MDR1 expression varies considerably across cancer types (*41*), with colon, renal and liver cancers exhibiting elevated MDR1 expression (*27, 28*). In contrast, other cancers such as lymphomas appear to have limited expression of MDR1 in cancer cell lines and patient tumors (*27, 28*), representing a potential patient population where PROTAC therapies may be more durable therapeutic outcomes. However, we demonstrated cancer cell lines that had non-detectable MDR1 protein levels induced MDR1 following chronic PROTAC exposure, acquiring resistance to PROTACs, suggesting the lack of MDR1 expression alone may not be sufficient to predict PROTACs response. MDR1 expression has been shown to be regulated by methylation, where many cancer cells display hypermethylation of the *ABCB1* promoter, maintaining gene suppression (*42*), thus, analysis of the methylation state of the *ABCB1* promoter in MDR1 non-expressing cells may be warranted to define a cancer patient population that may escape MDR1-mediated degrader-resistance. Further studies exploring the methylation state of the *ABCB1* promoter in cancer cells and its impact on degrader sensitivity, as well defining the methylation status of *ABCB1* in cancer cells that acquired resistance to degraders through upregulation of MDR1 will be of particular interest.

Small molecule inhibitors of MDR1 have been investigated in clinical trials as sensitizers to chemotherapies, however, these drugs have shown limited therapeutic benefit, with no MDR1 inhibitors FDA-approved for cancer therapy (*31*). MDR1 inhibitors have failed in clinic due to several limitations, such as poor drug accumulation and drug toxicities, prompting the search for alternative strategies to block MDR1-driven drug resistance (*32*). Several kinase inhibitors have been shown to directly inhibit MDR1 drug efflux activity, including a number of FDA-approved kinase inhibitors (*32, 43*). Here, we showed the FDA-approved MTOR inhibitor, RAD001, could be used to block MDR1 activity overcoming MDR1-mediated drug resistance in cancer cells. MTOR activation occurs frequently in cancers and targeting MTOR using RAD001 has been extensively tested in clinical trials, revealing RAD001 is safe, tolerable and has efficacy at blocking tumor growth in patients (*44*). RAD001 is currently used to treat several cancers, including renal cell carcinomas (RCC) (NCT00831480), which exhibits frequent overexpression of MDR1 (*45*). Further studies exploring whether RAD001 in combination with PROTACs targeting established drivers in RCC improves protein degradation and anti-tumor responses will be of interest. Moreover, exploring the impact of other dual MDR1/kinase inhibitors currently approved for cancer therapies, such as imatinib (*46*), or dasatinib (*47*), to improve PROTAC degrader efficiency and therapeutic responses may represent additional avenues to pursue for the treatment of MDR1 overexpressing cancers.

ErbB receptors are frequently altered in cancers, representing promising anti-cancer targets (*48*). Lapatinib is a highly selective EGFR, ERBB2 and ERBB4 inhibitor that is currently FDA-approved for the treatment of a variety of cancers (*49*). Notably, lapatinib has previously been shown to be a competitive inhibitor of MDR1 both *in vitro* and *in vivo (33)*, and our findings showed lapatinib could be used interchangeably with tariquidar to block or overcome MDR1-mediated resistance to PROTACs. Activation of ErbB receptors has been shown to promote resistance to KRAS^G12C^ or MEK inhibitors in colorectal cancers, where combination therapies of lapatinib and either KRAS^G12C^ or MEK inhibitors provided more durable therapies in tumor models (*36, 48*). Here, we demonstrated combining lapatinib with PROTACs targeting KRAS^G12C^ or MEK1/2 in MDR1-overexpressing CRC cells improved degradation KRAS^G12C^ or MEK1/2 and overall therapeutic responses. Our findings establish degradation of KRAS^G12C^ or MEK1/2 similarly induces ErbB3 activity and downstream AKT-signaling that is observed with small molecule inhibition, signifying blockade of compensatory ErbB3 signaling will also be required for KRAS^G12C^ or MEK1/2 degraders therapies to achieve durable response in CRC cells. ErbB receptor signaling has been shown to promote resistance to a variety of target agents including pan-Tyrosine Kinases (TK), AKT, RAF, MEK, and ERK inhibitors (*50*), and several PROTACs targeting these kinases have recently emerged. Determining whether lapatinib can globally improve degradation efficiency in combination with other PROTACs targeting K-ras effector pathways, as well as exploring lapatinib in combination with and KRAS^G12C^ or MEK1/2 degraders in other K-ras driven cancers such as lung and pancreatic cancers, will be of particular interest. Our preliminary *in vivo* studies suggest combining lapatinib and MEK degrader MS934 could have anti-tumor properties in K-ras mutant CRCs, however, more comprehensive *in vivo* studies exploring additional MDR1-overexpressing tumor models, as well as the potential cytotoxic effects of these combinations will be essential for therapeutic proof-of-concept.

## EXPERIMENTAL PROCEDURES

### Cell Lines

Cell lines were verified by IDEXX laboratories and free of mycoplasma. CAKI-1, DLD-1, HCT-15, HCT-116, NCI-H747, SW620, SW837, SW948, SW1116, and SW1463 cell lines were maintained in RPMI-1640 supplemented with 10% FBS, 100 U/ml Penicillin-Streptomycin and 2mM GlutaMAX. A1847, SUM159, and OVCAR3 cell lines were maintained in RPMI-1640 supplemented with 10% FBS, 100 U/ml Penicillin-Streptomycin, 2mM GlutaMAX, and 5 μg/mL insulin. LS513 and LS1034 cells were maintained in RPMI-1640 supplemented with 10% FBS, 100 U/ml Penicillin-Streptomycin, 2mM GlutaMAX, 1mM Sodium Pyruvate and 10mM HEPES. SKCO1 cells were maintained in MEM supplemented with 10% FBS, 100 U/ml Penicillin-Streptomycin, 2mM GlutaMAX and 1mM Sodium Pyruvate. PROTAC-resistant cells were maintained with 500nM PROTAC in the medium. All cells were kept at 37°C in a 5% CO_2_ incubator.

### Compounds

MEK1/2 degraders MS432 and MS934 were provided by the Jian Jin laboratory (*30*). All other compounds used are listed in Data File S2.

### Western Blotting

Samples were harvested in MIB lysis buffer (50 mM HEPES (pH 7.5), 0.5% Triton X-100, 150 mM NaCl, 1 mM EDTA, 1 mM EGTA, 10 mM sodium fluoride, 2.5 mM sodium orthovanadate, 1X protease inhibitor cocktail (Roche), and 1% each of phosphatase inhibitor cocktails 2 and 3 (Sigma)). Particulate was removed by centrifugation of lysates at 21,000 rpm for 15 minutes at 4°C. Lysates were subjected to SDS-PAGE chromatography and transferred to PVDF membranes before western blotting with primary antibodies. For a list of primary antibodies used, see (**Data File S2)**. Secondary HRP-anti-rabbit and HRP-anti-mouse were obtained from ThermoFisher Scientific. SuperSignal West Pico and Femto Chemiluminescent Substrates (Thermo) were used to visualize blots.

### Growth Assays

For short-term growth assays, 3000-5000 cells were plated per well in 96-well plates and allowed to adhere and equilibrate overnight. Drug was added the following morning and after 120 h of drug treatment, cell viability was assessed using the CellTiter-Glo Luminescent cell viability assay according to manufacturer (Promega). Students t tests were performed for statistical analyses and p values ≤ 0.05 were considered significant. For long term colony formation assays, cells were plated in 24-well dishes (1000-5000 cells per well) and incubated overnight before continuous drug treatment for 2 weeks, with drug and medium replenished twice weekly. Following the final treatment, cells were rinsed with PBS and fixed with chilled methanol for 10 min at −20°C. Methanol was removed by aspiration, and cells were stained with 0.5% crystal violet in 20% methanol for 1hr at room temperature.

### qRT-PCR

GeneJET RNA purification kit (Thermo Scientific) was used to isolate RNA from cells according to manufacturer’s instructions. qRT-PCR on diluted cDNA was performed with inventoried TaqMan® Gene Expression Assays on the Applied Biosystems 7500 Fast Real-Time PCR System. The TaqMan Gene Expression Assay probes (ThermoFisher Scientific) used to assess changes in gene expression include ABCB1 (Assay ID: Hs00184500_m1), and ACTB (control) (Cat # 4326315E).

### RNAi Knockdown Studies

siRNA transfections were performed using 25 nM siRNA duplex and the reverse transfection protocol. 3000-5000 cells per well were added to 96 well plates with media containing the siRNA and transfection reagent (Lipofectamine RNAiMax) according to the manufacturer’s instructions. Cells were allowed to grow for 120 h post-transfection prior to CellTiter Glo (Promega) analysis. Two-to-three independent experiments were performed with each cell line and siRNA. Students t tests were performed for statistical analyses and p values ≤0.05 were considered significant. For western blot studies, the same procedure was performed with volumes and cell numbers proportionally scaled to a 60mm or 10 cm dish, and cells were collected 72h post-transfection. siRNA product numbers and manufacturers are listed in (**Data File S2)**.

### Drug synergy analysis

Drug synergy was determined using SynergyFinder using the Bliss model and viability as the readout (https://doi.org/10.1093/nar/gkaa216). Each drug combination was tested in triplicate.

### Immunofluorescence

Cells were plated in a six-well plate with an 18-mm^2^ glass coverslip inside each well. Cells were fixed with 4% paraformaldehyde, permeabilized with 0.1% Triton X-100, blocked with 5% goat serum, and incubated with primary antibody (1:1000, anti-MDR1, Cell Signaling Technology) overnight at 4°C. The slides were washed with PBS and treated with secondary antibody (1:1000, FITC AffiniPure Donkey Anti-Rabbit IgG, Jackson Immunoresearch) for 1 hour at room temperature. Following antibody incubation, coverslips were mounted on slides using ProLong Gold Antifade Reagent with DAPI (4′,6-diamidino-2-phenylindole) (Thermo Fisher Scientific) and allowed to set overnight. Images were taken with a Nikon NI-U fluorescent microscope at 40x magnification.

### Rhodamine 123 Efflux Assay

Efflux assay was performed according to manufacturer’s protocol (Millipore Sigma #ECM910). Cells were resuspended in cold efflux buffer and incubated with Rhodamine 123 for 1 hr on ice. Cells were centrifuged and treated in warm efflux buffer with DMSO or drug for 30-60 min, washed with cold PBS, and effluxed dye was quantified with a plate reader at an excitation wavelength of 485 nm and an emission wavelength of 530 nm.

### Single Run Total Proteomics and Nano LC MS/MS

Parental or PROTAC-resistant cells were lysed in a buffer containing 50 mM HEPES pH 8.0 + 4% SDS, and 100 μg of protein was digested using LysC for 3 hours and trypsin overnight. Digested peptides were isolated using C-18 and PGC columns, then dried and cleaned with ethyl acetate. Three μg of proteolytic peptides were resuspended in 0.1% formic acid and separated with a Thermo Scientific RSLCnano Ultimate 3000 LC on a Thermo Scientific Easy-Spray C-18 PepMap 75μm x 50cm C-18 2 μm column. A 305 min gradient of 2-20% (180 min) 20%-28% (45 min) 28%-48% (20 min) acetonitrile with 0.1% formic acid was run at 300 nL/min at 50C. Eluted peptides were analyzed by Thermo Scientific Q Exactive or Q Exactive plus mass spectrometers utilizing a top 15 methodology in which the 15 most intense peptide precursor ions were subjected to fragmentation. The AGC for MS1 was set to 3×106 with a max injection time of 120 ms, the AGC for MS2 ions was set to 1×105 with a max injection time of 150 ms, and the dynamic exclusion was set to 90 s.

### Proteomics data processing

Raw data analysis of LFQ experiments was performed using MaxQuant software 1.6.0.1 and searched using Andromeda 1.5.6.0 against the Swiss-Prot human protein database (downloaded on April 24, 2019, 20402 entries). The search was set up for full tryptic peptides with a maximum of two missed cleavage sites. All settings were default and searched using acetylation of protein N-terminus and oxidized methionine as variable modifications. Carbamidomethylation of cysteine was set as fixed modification. The precursor mass tolerance threshold was set at 10 ppm and maximum fragment mass error was 0.02 Da. LFQ quantitation was performed using MaxQuant with the following parameters; LFQ minimum ratio count: Global parameters for protein quantitation were as follows: label minimum ratio count: 1, peptides used for quantitation: unique, only use modified proteins selected and with normalized average ratio estimation selected. Match between runs was employed for LFQ quantitation and the significance threshold of the ion score was calculated based on a false discovery rate of < 1%. MaxQuant normalized LFQ values were imported into Perseus software (1.6.2.3) and filtered in the following manner: Proteins identified by site only were removed, reverse, or potential contaminant were removed then filtered for proteins identified by >1 unique peptide. Protein LFQ values were log2 transformed, filtered for a minimum percent in runs (100%), annotated, and subjected to a Student’s *t*-test with comparing PROTAC-resistant cells *vs*. parental cells. Parameters for the Student’s *t*-test were the following: S0=2, side both using Permutation-based FDR <0.05. Volcano plots depicting differences in protein abundance were generated using R studio software and Prism graphics.

### Tumor xenograft experiment

Animal studies were conducted in accordance with the guidelines set forth by the Institutional Animal Care and Use Committee (Fox Chase Cancer Center IACUC # 16-16). 1 × 10^6^ LS513 cells were prepared in growth factor reduced Matrigel (Corning) 1:1 and injected into the right flank of 6- to 8- weeks old nude mice. Treatment with MS934 (50 mg/kg), Lapatinib (100mg/kg) or the combination (using the same dose as monotherapies) were started when tumors reached approximately 150 mm^3^ and maintained for two weeks. For *in vivo* studies, MS934 was resuspended in 5% N-methyl-2-pyrrolidinone (NMP), 5% Kolliphor HS-15 (Sigma) and 90% saline and delivered by intraperitoneal injection daily. Lapatinib was resuspended in 0.5% hydroxypropyl methylcellulose (Sigma) and 0.2% Tween-80 in distilled water pH 8.0. and delivered by oral gavage daily. Tumor volumes were evaluated every two days using a caliper and the volume was calculated applying the following formula: [(width)2 x (length)]/2.

## SUPPLEMENTAL INFORMATION

Supplemental information includes Supplemental Experimental Procedures, 7 figures and 2 Data Files.

## COMPETING INTERESTS

J.S.D. is an inventor on patent application WO2021026349A1 for using PROTACs in combination with dual MDR1 and kinase inhibitors for the treatment of cancer. J.J. and J. H. are inventors of a patent application filed by Icahn School of Medicine at Mount Sinai. The Jin laboratory received research funds from Celgene Corporation, Levo Therapeutics, and Cullgen Inc. J.J. is a cofounder, scientific advisory board member and equity shareholder in Cullgen Inc. and a consultant for Cullgen Inc., EpiCypher Inc., and Accent Therapeutics Inc. The other authors declare that they have no competing interests.

## AUTHOR CONTRIBUTIONS

J.S.D. wrote the manuscript. A.M.K. and S.M. performed CellTiter Glo assays and western blots. A.M.K., S.M., and D.A. performed colony formation assays. A.M.K. and J.S.D. performed all proteomics experiments and analysis. A.M.K. performed drug efflux assays, siRNA, qRT-PCR and immunofluorescence experiments. C.H.M. and D.A. performed xenograft studies. J.J. and J.H. provided MEK1/2 PROTACs MS432 and MS934. J.S.D. contributed to experimental design.

## ACKNOWLEDGMENTS

Funded by NIH CORE Grant CA06927 (Fox Chase Cancer Center), R01 CA211670 (J.S.D.), NIH T32 CA009035 (A.M.K), and Liz Tilberis Award Ovarian Cancer Research Alliance, 648813 (J.S.D).

## DATA and MATERIALS AVAILABILITY

Consortium through the PRIDE partner repository with the dataset identifier PXD029233. Reviewer account details: Username, Password: AtQ4UIA4.

## Notes

### Competing Interest Statement

J.S.D. is an inventor on patent application WO2021026349A1 for using PROTACs in combination with kinase inhibitors for the treatment of cancer. J.J. and J. H. are inventors of a patent application filed by Icahn School of Medicine at Mount Sinai. The Jin laboratory received research funds from Celgene Corporation, Levo Therapeutics, and Cullgen Inc. J.J. is a cofounder, scientific advisory board member and equity shareholder in Cullgen Inc. and a consultant for Cullgen Inc., EpiCypher Inc., and Accent Therapeutics Inc. The other authors declare that they have no competing interests.

