## Supplemental Figures and Legends for "MDR1 Drug Efflux Pump Promotes Intrinsic and Acquired Resistance to PROTACs in Cancer Cells"

### SUPPLEMENTAL FIGURES AND FIGURE LEGENDS

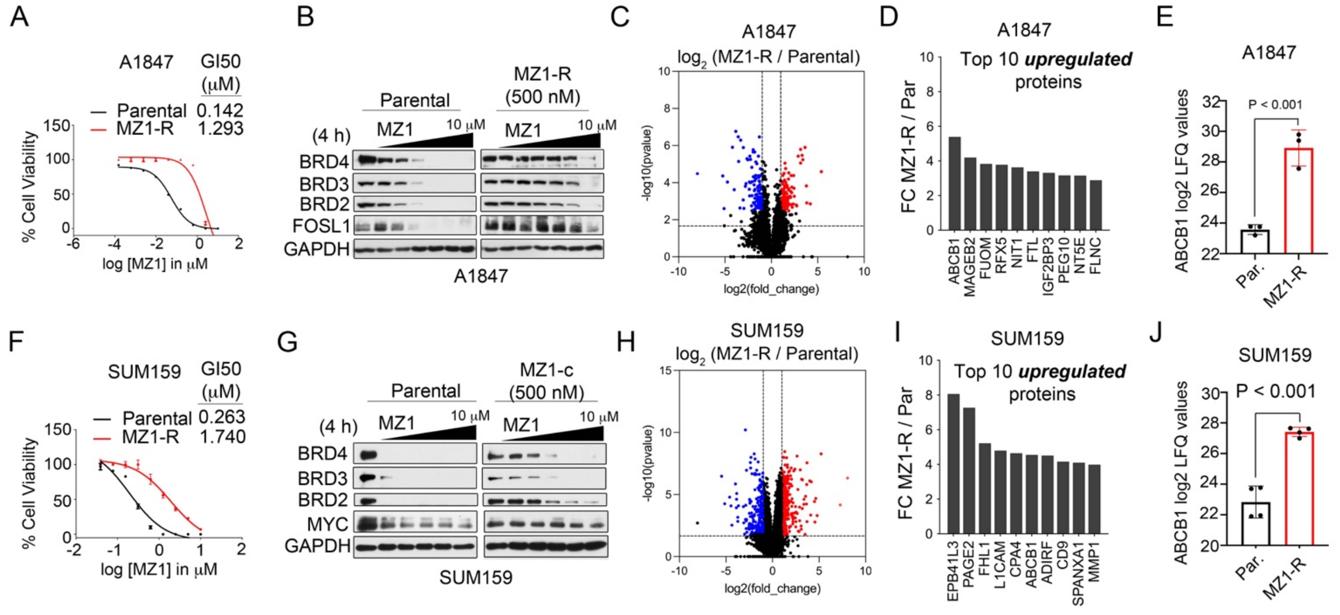


**Supplemental Figure 1. Proteomics Characterization of Degrader-Resistant Cancer Cell Lines**

(A) A1847 cells acquire resistance to MZ1. Parental or MZ1-resistant cells were treated with escalating doses of MZ1 for 5 d and cell viability assessed by CellTiter-Glo. MZ1-R treated cell viabilities normalized to DMSO treated degrader-R cells.

(B) Escalating doses of MZ1 less effective at inducing degradation of BET proteins in MZ1-R cells. A1847 parental or MZ1-R cells were treated with escalating doses of MZ1 (0, 0.039, 0.156, 0.625, 2.5 or 10 μM) for 24 h and degrader targets and downstream signaling determined by western blot. Blots are representative of 3 independent blots.

(C) Volcano plot depicts proteins elevated or reduced in MZ1-R relative to parental A1847 cells. Differences in protein log2 LFQ intensities amongst degrader-resistant and parental cells were determined by paired *t*-test Benjamini-Hochberg adjusted *P* values at FDR of <0.05 using Perseus software.

(D) Top 10 upregulated proteins in MZ1-R relative to parental A1847 cells.

(E) Bar graph depicts ABCB1 log2 LFQ values comparing MZ1-R relative to parental A1847 cells. Differences in ABCB1 log2 LFQ intensities amongst MZ1-R and parental cells were determined by paired *t*-test Benjamini-Hochberg adjusted *P* values at FDR of <0.05 using Perseus software.

(F) SUM159 cells acquire resistance to MZ1. Parental or MZ1-resistant cells were treated with escalating doses of MZ1 for 5 d and cell viability assessed by CellTiter-Glo. MZ1-R treated cell viabilities normalized to DMSO treated degrader-R cells.

(G) Escalating doses of MZ1 less effective at inducing degradation of BET proteins in MZ1-R cells. SUM159 parental or MZ1-R cells were treated with escalating doses of MZ1 (0, 0.039, 0.156, 0.625, 2.5 or 10 μM) for 24 h and degrader targets and downstream signaling determined by western blot. Blots are representative of 3 independent blots.

(H) Volcano plot depicts proteins elevated or reduced in MZ1-R relative to parental SUM159 cells. Differences in protein log2 LFQ intensities amongst degrader-resistant and parental cells were determined by paired *t*-test Benjamini-Hochberg adjusted *P* values at FDR of <0.05 using Perseus software.

(I) Top 10 upregulated proteins in MZ1-R relative to parental SUM159 cells.

(J) Bar graph depicts ABCB1 log2 LFQ values comparing MZ1-R relative to parental SUM159 cells. Differences in ABCB1 log2 LFQ intensities amongst MZ1-R and parental cells were determined by paired *t*-test Benjamini-Hochberg adjusted *P* values at FDR of <0.05 using Perseus software.

Data present in (A), (F) are triplicate experiments SD. *p ≤0.05 by student's t-test.

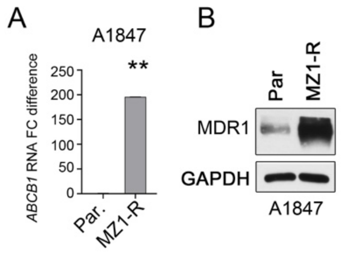


**Supplemental Figure 2**. **Chronic Exposure to Degraders Induces MDR1 Expression and Drug Efflux Activity**

(A) *ABCB1* mRNA levels are upregulated in MZ1-R cells relative to parental A1847 cells as determined by qRT-PCR.

(B) MDR1 protein levels are upregulated in MZ1-R cell lines relative to parental cells as determined by immunoblot. Blots are representative of 3 independent blots.

Data present in (A) are triplicate experiments SD. *p ≤0.05 by student's t-test.

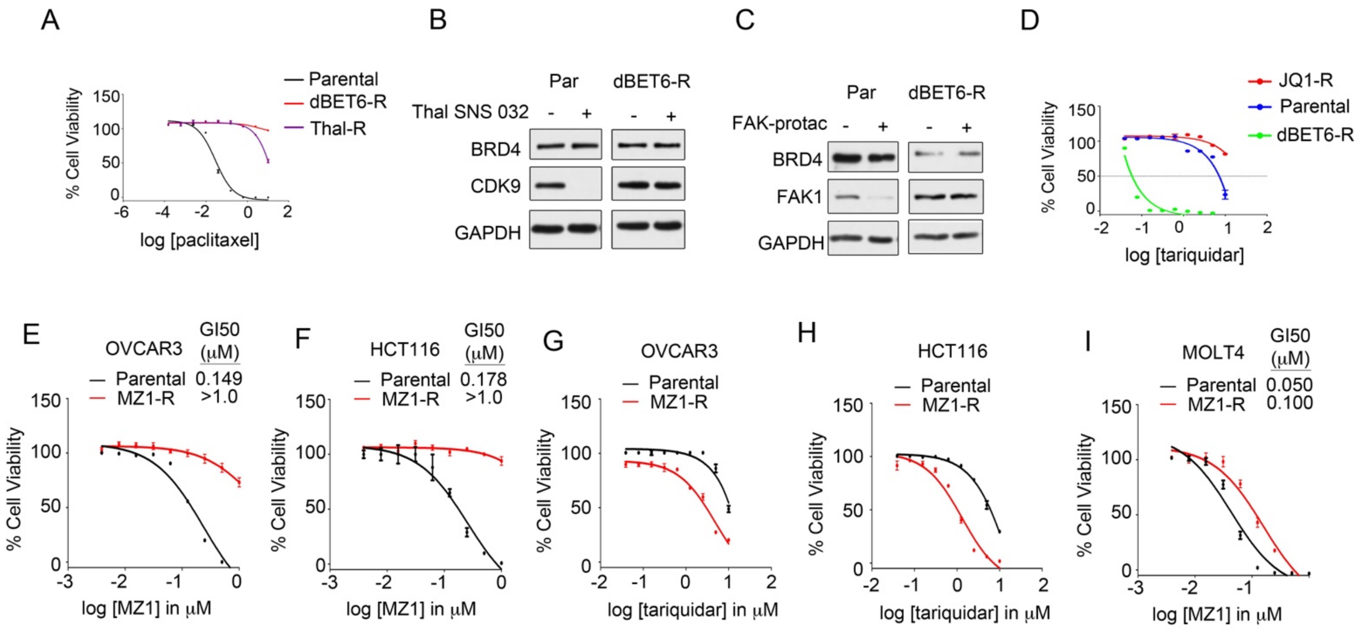


**Supplemental Figure 3. Blockade of MDR1 Activity Re-Sensitizes Degrader-Resistant Cells to PROTACs**

(A) Degrader-resistant cells also display resistance to MDR1-substrate paclitaxel. A1847 parental, dBET6-R or Thal-R cells were treated with doses of MZ1 for 5 d and cell viability assessed by CellTiter-Glo.

(B-C) Degrader-resistant cells cross-resistant to other PROTACs targeting different proteins. A1847 Parental or dBET6-R cells were treated with DMSO or (1 μM) Thal SNS 032 (B) or (1 μM) FAK-degrader-1 (C) for 24 h and proteins assessed by western blot.

(D) Chronic exposure to BET inhibitor JQ1 does not sensitize A1847 cells to MDR1 inhibition. A1847 parental, dBET6-R or JQ1-R cells were treated with doses of MZ1 for 5 d and cell viability assessed by CellTiter-Glo.

(E-F) OVCAR3 and HCT116 cells acquire resistance to MZ1. Parental or OVCAR3 (E) or HCT116 (F) MZ1-resistant cells were treated with escalating doses of MZ1 for 5 d and cell viability assessed by CellTiter-Glo. MZ1-R treated cell viabilities normalized to DMSO treated degrader-R cells.

(G-H) MZ1-resistant OVCAR3 and HCT116 cells exhibit increased sensitivity to MDR1 inhibitors. Cell-Titer Glo assay for cell viability of parental, MZ1-R OVCAR3 (G) or MZ1-R HCT116 (H) treated with increasing concentrations of MDR1 inhibitor tariquidar.

(I) MZ1 chronically exposed MOLT4 cells retain sensitivity towards MZ1 therapies. MOLT4 parental or MZ1-R cells were treated with doses of MZ1 for 5 d and cell viability assessed by CellTiter-Glo.

Data present in (A), (D), (E-I) are triplicate experiments SD. *p ≤0.05 by student's t-test.

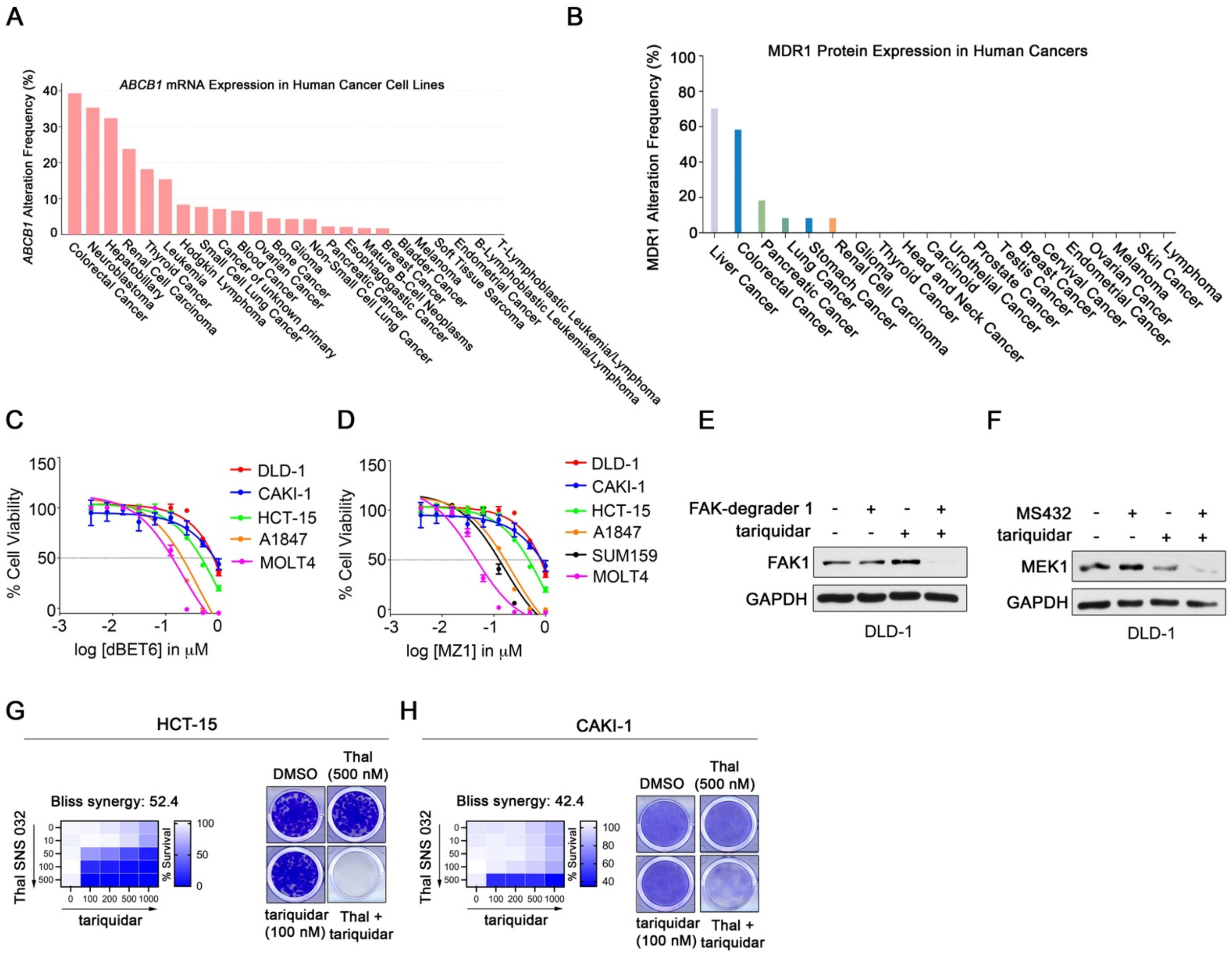


**Supplemental Figure 4. Overexpression of MDR1 Conveys Intrinsic Resistance to Degrader Therapies in Cancer Cells**

(A) Frequency of *ABCB1* mRNA overexpression across cancer cell line panel. Expression data was queried from cBioPortal for cancer genomics using Z-scores values of >2-fold-change for *ABCB1* mRNA levels (*1*).

(B) MDR1 protein expression across tumor samples by immunohistochemistry. MDR1 protein data was queried from the human protein atlas. MDR1 expression was determined by immunohistochemistry using MDR1 antibody CAB001716 (*2*).

(C-D) Cancer cells overexpressing MDR1 exhibit reduced sensitivity towards dBET6 or MZ1. Cancer cells were treated with escalating doses of dBET6 (C) or MZ1 (D) for 5 d and cell viability assessed by CellTiter-Glo. GI50 values were determined in Prism software.

(E-F) Inhibition of MDR1 sensitizes CRC cell line to other PROTACs. DLD-1 cells were treated with DMSO or (1 μM) FAK-degrader-1 (E) or (1 μM) MS432 (F) for 24 h and proteins assessed by western blot.

(G-H) Combining tariquidar with CDK9 degraders enhances growth inhibition of MDR1-overexpressing cell lines. Cell-Titer Glo assay for cell viability of HCT-15 (G) or CAKI-1 (H) cells treated with increasing concentrations of Thal SNS 032, tariquidar or the combination and bliss synergy scores determined. Cells were treated with DMSO, tariquidar (0.1 μM), Thal SNS 032 (5 μM) or the combination and colony formation assessed following 14-days of treatment. Colony formation image representative of 3 independent assays.

Data present in (C-D), (G-H) are triplicate experiments SD. *p ≤0.05 by student's t-test.

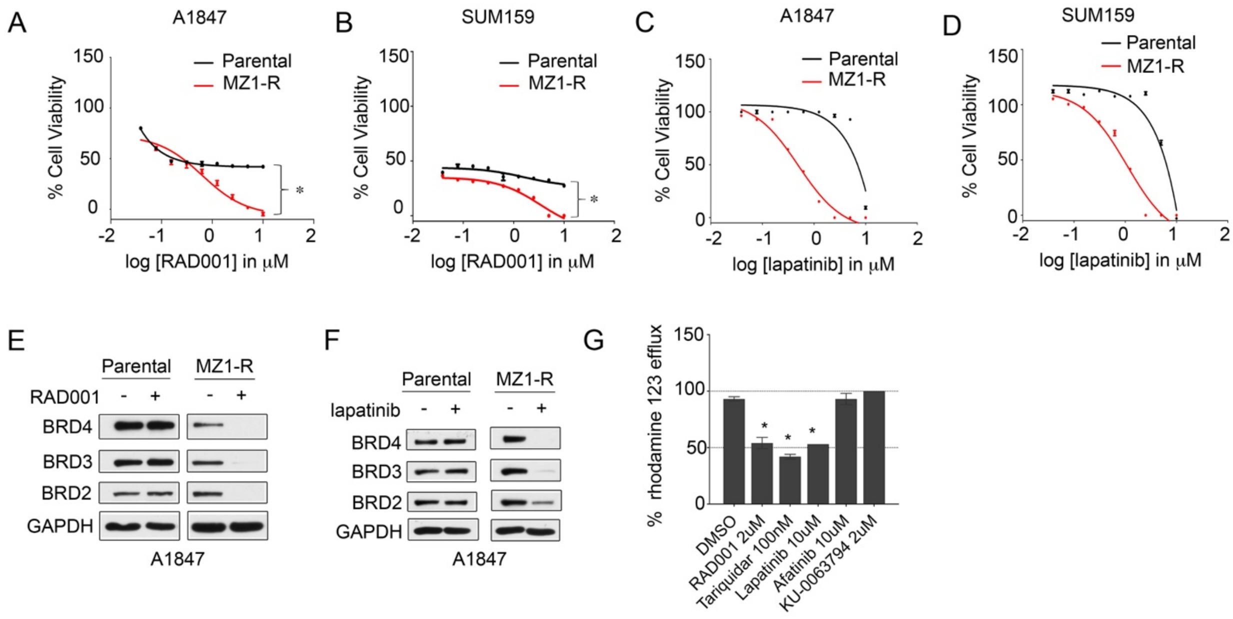


**Supplemental Figure 5. Re-Purposing Dual Kinase/MDR1 Inhibitors to Overcome Degrader Resistance in Cancer Cells**

(A-B) MZ1-resistant cells exhibit increased sensitivity towards RAD001. Cell-Titer Glo assay for cell viability of A1847 parental or MZ1-R (A) or SUM159 parental or MZ1-R (B) cells treated with increasing concentrations of RAD001.

(C-D) MZ1-resistant cells exhibit increased sensitivity towards lapatinib. Cell-Titer Glo assay for cell viability of A1847 parental or MZ1-R (A) or SUM159 parental or MZ1-R (B) cells treated with increasing concentrations of lapatinib.

(E-F) Treatment of MZ1-resistant cells with RAD001 or lapatinib promotes degradation of PROTAC-targets. A1847 parental, or MZ1-R cells treated with DMSO, RAD001 (2 μM) (E) or lapatinib (2 μM) (F) for 4 hours and proteins measured by western blot.

(G) Inhibitors Afatinib and KU-0063794 do not block MDR1 activity in degrader-resistant cells. Treatment of A1847 Thal-R cells with Afatinib or KU-0063794 does not reduce MDR1 drug efflux activity. A1847 Thal-R cells were treated with DMSO, 2 μM tariquidar, 2 μM RAD001, 2 μM lapatinib, 2 μM Afatinib or 2 μM KU-0063794 and Rhodamine 123 efflux assessed.

Data present in (A-D), (G) are triplicate experiments SD. *p ≤0.05 by student's t-test.

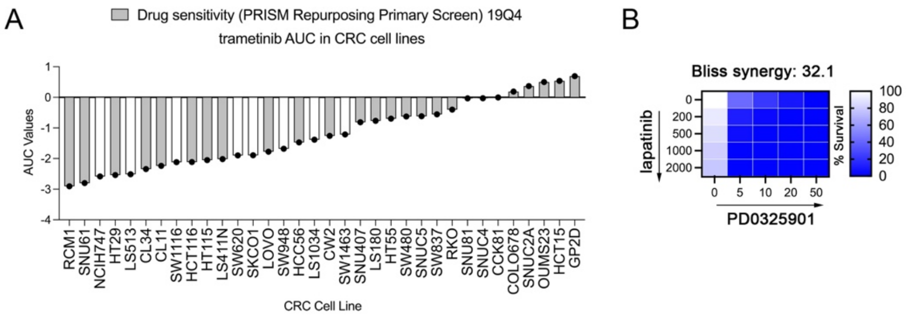


**Figure 6. Combining MEK1/2 Degraders with Lapatinib Synergize to Kill MDR1-Overexpressing K-ras Mutant CRC Cells**

(A) CRC cell lines explored in MS432 studies exhibit sensitivity towards MEK inhibition. Drug sensitivity profile of CRC cell lines to trametinib treatment. The bar graph depicts AUC from trametinib dose response studies. AUC data was queried from DepMap databases (*3*).

(B) Combining lapatinib with MEK inhibitors enhances growth inhibition of LS513 CRC cells. Cell-Titer Glo assay for cell viability of LS513 cells treated with increasing concentrations of lapatinib, PD0325901 or the combination and bliss synergy scores determined.

(C) Combining lapatinib with MEK degrader MS934 enhances growth inhibition of LS513 CRC cells. Cell-Titer Glo assay for cell viability of LS513 cells treated with increasing concentrations of lapatinib, MS934 or the combination and bliss synergy scores determined.

Data present in (B) and (C) are triplicate experiments SD. *p ≤0.05 by student’s t-test.

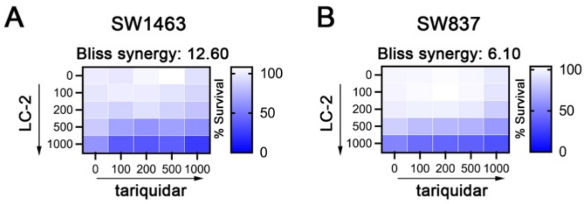


**Figure 7. Lapatinib-treatment improves KRAS$^{G12C}$ degrader therapies in MDR1-overexpressing CRC cell lines**

(A-B) Combining tariquidar with KRAS$^{G12C}$ degraders display less drug synergy than when combined with lapatinib in CRC cells. Cell-Titer Glo assay for cell viability of SW1463 (A) or SW837 (B) cells treated with increasing concentrations of lapatinib, LC-2 or the combination and bliss synergy scores determined.

Data present in (A-B) are triplicate experiments SD. *p ≤0.05 by student's t-test.

### REFERENCES


1. J. Gao, B. A. Aksoy, U. Dogrusoz, G. Dresdner, B. Gross, S. O. Sumer, Y. Sun, A. Jacobsen, R. Sinha, E. Larsson, E. Cerami, C. Sander, N. Schultz, Integrative Analysis of Complex Cancer Genomics and Clinical Profiles Using the cBioPortal. *Science signaling* **6**, pl1-pl1 (2013); published online Epub04/02 (10.1126/scisignal.2004088).
2. M. Uhlén, L. Fagerberg, B. M. Hallström, C. Lindskog, P. Oksvold, A. Mardinoglu, Å. Sivertsson, C. Kampf, E. Sjöstedt, A. Asplund, I. Olsson, K. Edlund, E. Lundberg, S. Navani, C. A. Szigyarto, J. Odeberg, D. Djureinovic, J. O. Takanen, S. Hober, T. Alm, P. H. Edqvist, H. Berling, H. Tegel, J. Mulder, J. Rockberg, P. Nilsson, J. M. Schwenk, M. Hamsten, K. von Feilitzen, M. Forsberg, L. Persson, F. Johansson, M. Zwahlen, G. von Heijne, J. Nielsen, F. Pontén, Proteomics. Tissue-based map of the human proteome. *Science* **347**, 1260419 (2015); published online EpubJan 23 (10.1126/science.1260419).
3. J. Barretina, G. Caponigro, N. Stransky, K. Venkatesan, A. A. Margolin, S. Kim, C. J. Wilson, J. Lehár, G. V. Kryukov, D. Sonkin, A. Reddy, M. Liu, L. Murray, M. F. Berger, J. E. Monahan, P. Morais, J. Meltzer, A. Korejwa, J. Jané-Valbuena, F. A. Mapa, J. Thibault, E. Bric-Furlong, P. Raman, A. Shipway, I. H. Engels, J. Cheng, G. K. Yu, J. Yu, P. Aspesi, M. de Silva, K. Jagtap, M. D. Jones, L. Wang, C. Hatton, E. Palescandolo, S. Gupta, S. Mahan, C. Sougnez, R. C. Onofrio, T. Liefeld, L. MacConaill, W. Winckler, M. Reich, N. Li, J. P. Mesirov, S. B. Gabriel, G. Getz, K. Ardlie, V. Chan, V. E. Myer, B. L. Weber, J. Porter, M. Warmuth, P. Finan, J. L. Harris, M. Meyerson, T. R. Golub, M. P. Morrissey, W. R. Sellers, R. Schlegel, L. A. Garraway, The Cancer Cell Line Encyclopedia enables predictive modeling of anticancer drug sensitivity. *Nature* **483**, 603-607 (2012); published online Epub03/28 (10.1038/nature11003).
